# Systematic fMRI signal differences across cohorts alter lifespan trajectories of functional brain networks

**DOI:** 10.64898/2026.01.15.699580

**Authors:** Micaela Y. Chan, Liang Han, Gagan S. Wig

## Abstract

Large-scale lifespan neuroimaging studies increasingly integrate data across distinct cohorts to characterize trajectories of brain development and aging. However, systematic differences in acquisition protocols and hardware across cohorts can alter signal characteristics in ways that bias downstream analyses. Here, we examine three cohorts from the Human Connectome Project (HCP), spanning development (HCP-D), young adulthood (HCP-YA) and aging (HCP-A), to illustrate this issue and evaluate existing strategies to mitigate it. HCP has set standards for open, deeply phenotyped, high-resolution human neuroimaging, which are frequently used as high-quality reference datasets in tool validation, replication studies, and cross-cohort meta-analyses. However, neuroimaging acquisitions have differed across HCP cohorts because of changes in scanner hardware and acquisition sequences across study phases. Because of HCP’s widespread usage, even modest protocol differences between cohorts–and their downstream effects–can have outsized impacts on the field of neuroscience research. Our analysis reveals that the HCP-YA cohort exhibits systematically weaker temporal signal-to-noise ratio (tSNR) relative to HCP-D/A. These signal quality discrepancies propagate to downstream analyses, leading to differences in overall resting-state functional correlations and whole-brain and node-level measures of resting-state network organization (e.g., system segregation, modularity, participation coefficient). Consistent with protocol-driven signal differences, resting-state network measures derived from HCP-YA depart from expected lifespan trajectories, as confirmed by examination of two other lifespan datasets. Harmonization approaches accounting for protocol and scanner-model differences substantially lessen these artifactual differences in brain network measures. Our findings underscore that signal differences do not merely introduce noise, but can qualitatively alter estimated lifespan trajectories of functional network organization, including partially inverting expected lifespan patterns. Without appropriate harmonization, analyses that combine HCP cohorts can therefore result in biologically misleading inferences about brain development and aging. We demonstrate how small acquisition differences bias resting-state-derived network metrics, and how these effects can be mitigated. This work advances best practices for valid inference in multi-cohort lifespan neuroscience research.

## 1. INTRODUCTION

A growing interest in understanding brain changes across the lifespan has led to an increased number of large multi-site studies and integrative approaches that combine multiple studies which have acquired data across distinct life course segments (Bethlehem et al., 2022; Sun et al., 2025; Zhou et al., 2023). While these efforts have allowed researchers to gain a comprehensive understanding of changes and differences in brain structure and function across broad segments of the human lifespan, it is critical to ensure data across studies are comparable and harmonized in ways that do not confound interpretations of the derived measures (Pomponio et al., 2020). Many recently published studies using multi-site data have employed some form of harmonization or control for site-differences in their models. However, this approach often places the primary focus on “site” (referring to the physical location of data collection) as the source of variability, rather than explicitly addressing protocol or hardware differences, which may be equally, if not more important to control for. For instance, even when efforts are made to keep imaging protocols as similar as possible, unanticipated hardware changes (e.g., scanner upgrade/decommissioning) and the necessity of using specific protocols to accommodate special populations (e.g., children, older adults, patients) can introduce systematic and confounding variations into comparisons.

Relevant to the above issues, our report focuses on a collection of valuable and readily available datasets that, when combined, have enabled novel discoveries of brain structure and function across the lifespan: the Human Connectome Project (HCP), HCP-Development, and HCP-Aging (Bookheimer et al., 2019; Glasser et al., 2013; Somerville et al., 2018). These multi-site data collection efforts were designed to be combined despite having been collected on different scanners and with different protocols due to hardware upgrades. Here, we demonstrate the necessity of proper data harmonization to prevent misinterpretation of downstream results.

The HCP and its ancillary studies have established themselves as invaluable resources for exploring questions about brain development and aging in health and disease. The scale and richness of these datasets, coupled with rigorous quality control and the HCP team’s commitment to releasing both raw and fully processed data, represent an exceptional contribution to the field. With the release of the newly processed original HCP data (*Aging Adult Brain Connectome Release 1*; *HCP-Young Adult 2025 Release*), it is clear that the team continues to enhance data usability and reproducibility, making these resources uniquely powerful for lifespan neuroscience. However, it is important to acknowledge that due to scanning protocol and hardware changes, there exist distinct differences across the efforts. The HCP-YA cohort was collected on a (*i*) different scanner and with (*ii*) different protocols than those later used for the HCP-D/A datasets. Some of these changes were made to achieve the goal for HCP-D and HCP-A to have a protocol that is usable across multiple sites and a much wider age-span (e.g., standard scanner available at multiple locations, shorter scanning time for children and elderly; Bookheimer et al., 2019; Harms et al., 2018). At the time, rigorous efforts were made to ensure that newer protocols in the developmental and aging cohort were as similar as possible to the original young adult cohort collected on the connectome scanner (see Harms et al., 2018 for comparison of protocols and signals). Additionally, even within the same acquisition protocol (e.g., voxel size, repetition time), fMRI temporal signal-to-noise-ratio (tSNR) can be influenced by multiple factors such as hardware (e.g., scanner model, scanner field strength, head coil channel) and participant trait characteristics (e.g., motion; Van Dijk et al., 2012). For example, in the case of comparing the original HCP-YA cohort with HCP-D or HCP-A, the resting-state protocol and scanner hardware were different: HCP-YA collected resting-state scan with a 720ms repetition time (TR; LR/RL phase encoding) from a custom Siemens Skyra scanner, and HCP-D/A collected resting-state scan with a 800ms TR (AP/PA phase encoding) from Siemens Prisma scanners. These protocol and hardware differences can alter signal characteristics (see Adhikari et al., 2019 for influence of scan parameters on signal quality), with downstream effects on resting-state network measures derived from the scans, potentially obscuring true neurobiological differences across the lifespan.

Our focus is on the functional connectome derived from measures of resting-state functional MRI, given the increasing recognition of its role in supporting distributed and coordinated brain activity that underlies cognition and behavior. There are many ways to analyze resting-state signals and their coordinated patterns of organization, but here we interrogate several commonly examined measures of resting-state correlations that have been examined in the context of lifespan differences and changes (e.g., Betzel et al., 2014; Chan et al., 2014; Geerligs et al., 2015; Tooley et al., 2020; Winter-Nelson et al., 2026). Specifically, we examine the mean level of resting-state functional correlation (RSFC) across the whole brain, within and between known functional systems, and four widely used graph-theoretical network measures: system segregation (Chan et al., 2014), modularity (Newman, 2004b), participation coefficient (Guimerà & Amaral, 2005) and clustering coefficient (Watts & Strogatz, 1998). No a priori assumption is made as to whether certain measures are more susceptible to acquisition-related differences, instead, we aim to evaluate the extent and generality to which these measures of large-scale brain network organization derived from RSFC are affected by acquisition-differences.

In the present work, we are not aiming to pinpoint whether the protocol or hardware drives the cohort-level differences across the HCP datasets, as the two are completely intertwined, but to show that notable differences in signal quality, as quantified by tSNR, and their effect on downstream resting-state network measures exist. tSNR is examined because it is commonly used to evaluate signal quality (Parrish et al., 2000; Welvaert & Rosseel, 2013) and summarizes the aggregated influence of acquisition parameters and hardware (e.g., TR, phase-encoding direction; Jamil et al., 2021; Krüger & Glover, 2001; Triantafyllou et al., 2005). The goal is to help researchers wanting to combine these data to interrogate lifespan questions become aware of this issue and properly harmonize their data.

Importantly, to demonstrate that the observed cohort-related effects reflect acquisition differences rather than general properties of lifespan functional organization, we additionally evaluate two independent lifespan datasets as reference cohorts: the Nathan Kline Institute Rockland Sample (NKI; Nooner et al. 2012) and the Dallas Lifespan Brain Study (DLBS; Park et al. 2025). These datasets differ in node definitions and preprocessing strategies, providing a stringent test of whether any observed effects generalize beyond specific analytic choices.

To summarize, the present study investigates how subtle differences in HCP acquisition protocols result in observable differences in fMRI tSNR when combining these three valuable datasets to examine age-related questions, and that these differences introduce systematic confounds in resting-state derived network measures if not adequately corrected. Across analyses, we show that unharmonized data from HCP cohorts can partially invert expected developmental and aging trajectories of large-scale network organization.

## 2. METHODS

### 2.1 Participants

The study used data from three publicly available datasets from the Human Connectome Project (HCP): the HCP-Young Adult (HCP-YA), HCP-Development (HCP-D), and HCP-Aging (HCP-A) cohorts (Bookheimer et al., 2019; Glasser et al., 2013; Somerville et al., 2018); the Nathan Kline Institute Rockland Sample (NKI; Nooner et al. 2012); and the Dallas Lifespan Brain Study (DLBS; Park et al. 2025).

All datasets used in this study were obtained from publicly available, de-identified datasets collected with appropriate ethical approval and informed consent. The present analyses involved secondary use of these data and were approved by the institutional review board (IRB) at the University of Texas at Dallas.

#### 2.1.1 Human Connectome Project (HCP)

For all three HCP cohorts, the scanning protocol was approved by the respective review boards of the data collection sites. HCP-YA’s 3T MRI data were collected at Washington University in St. Louis. HCP-D data were acquired at the University of California-Los Angeles, the University of Minnesota, Washington University in St. Louis and Harvard University. For HCP-A, Massachusetts General Hospital was used as the 4^th^ data collection site instead of Harvard. All participants or their legal guardian provided written informed consent.

##### HCP-Young Adult (HCP-YA)

The HCP-YA participants were drawn from the HCP S1200 release dataset (downloaded in March 2023 from https://www.humanconnectome.org; n=1206; age range: 21–37y; Glasser et al. 2013; Marcus et al. 2013). The final sample used in the current study included participants with at least 10 minutes of clean resting-state data per session (see Resting-state fMRI preprocessing), mean head motion below filtered framewise displacement (fFD; Gratton et al., 2020) of 0.04, average tSNR across run was not below 3SD of the HCP-YA cohort’s mean tSNR, and had no QC-issue reported by HCP raters (n=764 [F=422], M_age_=28.6, SD_age_=3.7).

##### HCP-Development (HCP-D)

The HCP-D data were drawn from the HCP-D release 2.0 (https://nda.nih.gov; n=625, age range: 5–21). The final sample included participants with at least 10 minutes of clean resting-state data, mean head motion (fFD) < 0.04, and the average tSNR across runs was not below 3SD of the HCP-D cohort (see Resting-state fMRI preprocessing; n=519 [F=288], M_age_=15.1, SD_age_=3.9).

##### HCP-Aging (HCP-A)

The HCP-A data were drawn from the HCP-A release 2.0 (https://nda.nih.gov; n=725, age range: 36–100y). The final sample included participants with at least 10 minutes of clean resting-state data (see Resting-state fMRI preprocessing), aged 90 or below, mean head motion (fFD) < 0.04, and the average tSNR across runs was not below 3SD of the HCP-A cohort (n=531 [F=299], M_age_=58.7, SD_age_=14.8). Nine participants above the age of 90 were available; however, because ages above 90 were jittered to 100 in the public release to protect participant privacy, these extreme age values skewed the age trend. Accordingly, participants above the age of 90 were excluded from the analysis.

#### 2.1.2 Nathan Kline Institute Rockland Sample (NKI)

The Nathan Kline Institute – Rockland Sample (NKI) is an open-access neuroimaging dataset that spans a large age range from childhood to older age (Nooner et al., 2012). All participants provided written consent, and study procedures were approved by the Nathan Kline Institute for Psychiatric Research Institutional Review Board. For minors, both written assent from the participant and written consent from a parent or legal guardian were obtained. The present study included 618 individuals (age range: 6–83 years; M_age_=40.6, SD_age_=20.4) who completed both anatomical and resting-state fMRI and passed fMRI motion processing criteria (see below).

#### 2.1.3 Dallas Lifespan Brain Study (DLBS)

Participants were recruited from the Dallas–Fort Worth community as part of the Dallas Lifespan Brain Study (DLBS), which examined healthy aging and cognition (Park et al., 2025). All participants provided written consent, and study procedures were approved by the Institutional Review Boards at the University of Texas at Dallas and UT Southwestern Medical Center. The present study included 358 adults (age range: 20– 89 years; M_age_=56.9, SD_age_=18.4) who completed both anatomical and resting-state fMRI and passed fMRI motion processing criteria (see below).

### 2.2 Neuroimaging

#### 2.2.1 Structural imaging acquisition and preprocessing

##### HCP

Each participant underwent T1-weighted (T1w) magnetization-prepared rapid acquisition gradient echo (MP-RAGE) scans (TR=2400ms, TE=2.14ms, TI=1000ms, flip angle=8°, 0.7mm isotropic voxels, 256 slices, FOV=224×224mm) and T2-weighted (T2w) scans (TR=3200ms; TE=565ms; 0.7mm isotropic voxels, 256 slices, FOV=224×224mm).

The raw T1w and T2w images were locally processed using the HCP-pipeline (v4.2), including the pre-FreeSurfer, FreeSurfer, and post-FreeSurfer structural processing steps. FreeSurfer 6.0., FSL 6.0.4, and MATLAB 2019b were used as part of the HCP-pipeline.

##### NKI

T1w MP-RAGE structural images were collected with the following parameters: TR=2500ms, TE=3.5ms, TI=1200ms, flip-angle=8°, 1mm isotropic voxels, 192 slices, FOV=256×256 mm. The structural images were not downloaded or further processed because the functional data were already registered to standard MNI space.

##### DLBS

T1w MP-RAGE structural images were collected with the following parameters: TR=8.1ms, TE=3.7ms, flip-angle=12°, 1mm isotropic voxels, 160 slices, FOV=204×256 mm. Data were processed with FreeSurfer v5.3, with manual editing to ensure proper surface estimation (see Savalia et al., 2017 for details).

#### 2.2.2 Resting-state fMRI acquisition

##### HCP

Eyes-opened with fixation (white crosshair, black background) resting-state functional MRI were acquired from each participant using a gradient-echo EPI sequence. Parameters for HCP-YA and HCP-D/A differed slightly.

###### HCP-YA

RL/LR phase encoding, multiband factor=8, TR=720ms, TE=33.1ms, flip-angle=52°, 2mm isotropic voxels, 72 slices, FOV=208×180mm, 1200 volumes (∼14.4 min). HCP-YA participants had a total of 4 runs, totaling ∼58 min of resting-state data.

###### HCP-D/A

AP/PA phase encoding, multiband factor=8, TR=800ms, TE=37ms, flip-angle=52°, 2mm isotropic voxels, 72 slices, FOV=208×180mm, 488 volumes (∼6.5 min). HCP-D/A participants aged 8 or above had a total of 4 runs, totaling ∼26 min of resting-state data. HCP-D participants below the age of 8 had 6 runs of the shorter scans (∼3.4 min), totaling ∼20 min of resting-state data.

##### NKI

Eyes-opened resting-state functional MRI without a fixation cross were acquired from each participant. Three different sequences were collected, the main text included analyses using the sequence with parameters closest to HCP-style data (shortest TR, multiband), but the supplemental information includes the other two sequences (see **Supplementary Information (SI); SI sections are ordered based on their appearance in the Results section**). The sequence used in the main analysis had the following parameters: multiband factor=4, TR=645ms, TE=30ms, flip-angle=60°, 3mm isotropic voxels, 40 slices, FOV=200×200mm, 900 volumes (∼9.7min).

##### DLBS

Eyes-opened with fixation (white crosshair, black background) resting-state fMRI were acquired from each participant: TR=2000ms, TE=25ms, flip-angle=80°, 3.5mm isotropic voxels, 43 slices, FOV=220×220 mm. Participants with data collected before the year 2011 had a single run of resting-state fMRI with 154 volumes (∼5.1 min), whereas participants collected after had two runs with 180 volumes each (12 min). Of the participants who passed quality control, 223 participants had 5 minutes of data, and 135 participants had 12 minutes of data.

#### 2.2.3 Resting-state fMRI preprocessing

##### HCP

Resting-state fMRI data used in the present study were locally processed using the HCP-pipeline (v4.2) fMRI volume pipeline, motion-scrubbing, and the HCP-pipeline fMRI surface pipeline (v4.2). **SI 1.1** includes the results using the downloaded “Preprocessed Recommended fMRI Data” from HCP-YA (connectome-DB) and HCP-D/A (NDA version 2.0).

After the HCP-pipeline for anatomical data was completed, HCP-pipeline fMRI volumetric pipeline was used to correct for gradient-nonlinearity-induced distortion, realignment of time series, and EPI distortion correction. The intensity of the processed volumetric BOLD data was scaled to a mode of 1000 (Ojemann et al. 1997).

Additional resting-state functional correlation (RSFC) specific processing and head motion correction were applied to reduce spurious variance from non-neuronal activity (Power et al. 2014): (i) demeaning and detrending; (ii) multiple regression to remove variance associated with global brain signal (GSR), ventricular signal, white matter signal, their derivatives, and the “Friston24” motion regressors; (iii) “Scrubbing”, which involves identifying and interpolating motion contaminated volumes defined by a fFD (Gratton et al., 2020) threshold greater than 0.04mm, as well as any volumes between two contaminated volumes fewer than 5 volumes apart (Power et al., 2014); (iv) bandpass filtering (0.009-0.08 Hz); (v) removing the interpolated volumes that were temporarily retained during bandpass filtering to preserve temporal continuity. We note that the interpolated volumes were temporarily introduced to avoid artifacts during filtering (i.e., to preserve temporal continuity and minimize edge artifacts during filtering) (Power et al., 2014). However, these interpolated volumes were not true neural signals, therefore, they were excluded from functional connectivity calculations to prevent introducing artificially created data that could bias correlation estimates. The HCP-pipeline surface processing was used to register the RSFC/motion processed data to the fs_LR (32k) surface-based atlas for analysis (Van Essen et al. 2012).

Functional connectivity estimates are sensitive to the amount of data included (Birn et al., 2013; Laumann et al., 2015; Noble et al., 2017), and differences in data quantity across participants biases comparison of measures of network organization derived from functional connectivity (Han et al., 2024). Therefore, the number of volumes in each session after motion “scrubbing” was fixed at 10 minutes (750 volumes for HCP-D/A, 834 volumes for HCP-YA), which ensured comparability across individuals within and across HCP cohorts.

Although many participants had longer acquisitions, the 10-minute criterion was selected to balance equating the amount of usable data across participants and maximizing participant retention within each cohort. Specifically, the youngest participants in HCP-D below the age of 8y had shorter resting-state scans (∼20min of acquisition), resulting in less usable data after motion censoring. The selected data-length criterion corresponded to approximately 50% of the initially available data in this lowest-duration group. The patterns observed across cohorts were consistent when a more stringent data-length criterion was applied (see **SI 1.8** showing qualitatively similar patterns across HCP cohorts using 20 minutes of data).

##### NKI

The processed NKI data were downloaded from the S3 bucket in May 2025. The 10-minute resting-state sequence with a 645ms TR was used in the primary analysis (see **SI 1.7** for similar relations between age and tSNR/network measures using the two other resting-state sequences [TR = 1400ms, TR = 2500ms]).

The processing steps were performed with the Configurable Pipeline for the Analysis of Connectomes (C-PAC), a Nipype-based, open-source pipeline for functional MRI data. Native-space functional images were slice-timing corrected, coregistered to MNI space using ANTs, intensity-normalized, and brain-masked using FSL. RSFC processing included polynomial detrending; nuisance regression of mean CSF, aCompCor components from white matter (5 components), and global signal; as well as the Friston 24 motion regressors.

While scrubbing was not used on the downloaded output, to match the processing of the other two datasets, volumes with motion above 0.3mm FD were discarded. To ensure comparability within the NKI dataset and to minimize bias in functional connectivity estimates due to unequal data length, each individual’s time series was standardized to the same length. Each session was standardized to 466 volumes, equivalent to ∼5 minutes, where the final data length corresponded to approximately 50% of the initially available data. Participants with less than 466 volumes of data after removing motion-contaminated volumes were not included.

##### DLBS

The DLBS resting-state fMRI data used in the present study were locally processed (see Chan et al., 2018 for details). Preprocessing included slice-timing correction, realignment, and RSFC processing identical to that applied to the locally processed HCP data, but with a non-filtered framewise displacement threshold of 0.3 mm. Workbench v1.4.2 was used to register the RSFC- and motion-processed data to the fs_LR (32k) surface-based atlas for analysis (Van Essen et al., 2012). To maintain consistency in functional connectivity estimation across DLBS participants, each session was standardized to 75 volumes, equivalent to 2.5 minutes. This data-length criterion has been used in prior work with this dataset (Chan et al., 2018) and corresponds to approximately 50% of the initially available data in participants with the shortest scans.

Because NKI and DLBS have shorter scan durations, the resulting data lengths were shorter than those used for the HCP cohorts. Although shorter time series reduce test-retest reliability (Birn et al., 2013; Laumann et al., 2015; Noble et al., 2017), standardizing data length within each dataset (Han et al., 2024) ensured unbiased comparisons across participants within each dataset. In the current work, cross-dataset comparisons were interpreted in terms of relative patterns rather than absolute values.

### 2.3 Brain network construction

#### HCP & DLBS

For each session of surface-mapped resting-state data, a functional correlation matrix was generated using a 441-node atlas (Chan et al., 2014). These nodes were defined based on boundary-based analyses (Chan et al., 2014; Wig et al., 2014), and labeled according to their spatial overlap with a published vertex-wise community map (Power et al., 2011). The BOLD time series of all vertices within each node were averaged to obtain the node’s mean time series. A correlation matrix (brain network) was constructed by computing the pairwise Fisher’s z-transformed Pearson correlation between nodes. In HCP data, one node was excluded due to its location overlapping with the more liberal medial wall used by the HCP-pipeline. Thus, a 440 × 440 matrix was available for each HCP participant. DLBS used the same 441 surface-based nodes, but because the preprocessing pipeline employed a more conservative medial wall mask than HCP, all 441 nodes were retained.

#### NKI

For NKI, a 264-node atlas (Power et al., 2011) was used to generate the RSFC matrix. This volumetric node set was used to stay consistent with the publicly available processed outputs. This differed from the surface-based parcellations used for the HCP and DLBS datasets, and was intentionally retained to provide a test of whether observed effects generalize across commonly used node definitions and atlases rather than being specific to a particular data processing stream. These nodes were defined in volumetric (MNI) space, and the BOLD signal from all voxels within each node was averaged to obtain a mean time series for each node. Pairwise Fisher’s z-transformed Pearson’s correlations were then computed between all nodes to form a 264 × 264 matrix for each participant.

### 2.4 Measures

#### 2.4.1 Temporal signal-to-noise ratio (SNR)

Temporal signal-to-noise ratio (tSNR) provides an estimate of the reliability of the BOLD signal over time within each voxel. It is defined as:

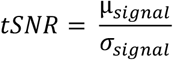

where µ_*signal*_ is the mean BOLD signal intensity over time for a given voxel, and σ_*signal*_ is the standard deviation of the BOLD signal intensity of the voxel’s time series. Higher tSNR values indicate that the BOLD signal is relatively stable (i.e., large mean signal relative to temporal fluctuations), whereas lower values reflect greater instability in the time series (possibly due to noise contributions).

For each participant and each resting-state run, voxel-wise tSNR was calculated after preprocessing and registration to MNI space. In order to capture signal properties that can be largely attributed to the acquisition (e.g., scanner, protocol, and physiological noise structure) rather than effects introduced by downstream preprocessing (e.g., spatial smoothing, temporal filtering, nuisance regression, motion correction), tSNR was calculated from processed time series of each run prior to volume censoring and surface mapping. These tSNR estimates were subsequently used to characterize individual and cohort-level differences in signal quality and to evaluate how such differences may propagate to downstream functional connectivity and network measures (also see **SI 1.1, Fig. S1** showing consistent patterns of cohort-level differences when tSNR was extracted from “Preprocessed Recommended fMRI Data” provided by the HCP team, and **SI 1.2, Fig. S2B** showing results from when tSNR was extracted from data with high-motion volumes removed).

The mean tSNR for a run was then obtained by averaging across all voxels within a whole-brain mask. Cohort-level tSNR values were derived by averaging across all participants within a cohort (used in **Fig. 1A & 1B**). Participant-level tSNR was defined as the mean tSNR for a given participant, computed from a single run or averaged across multiple runs when available (used in **Fig. 1C**).

**Figure 1.**
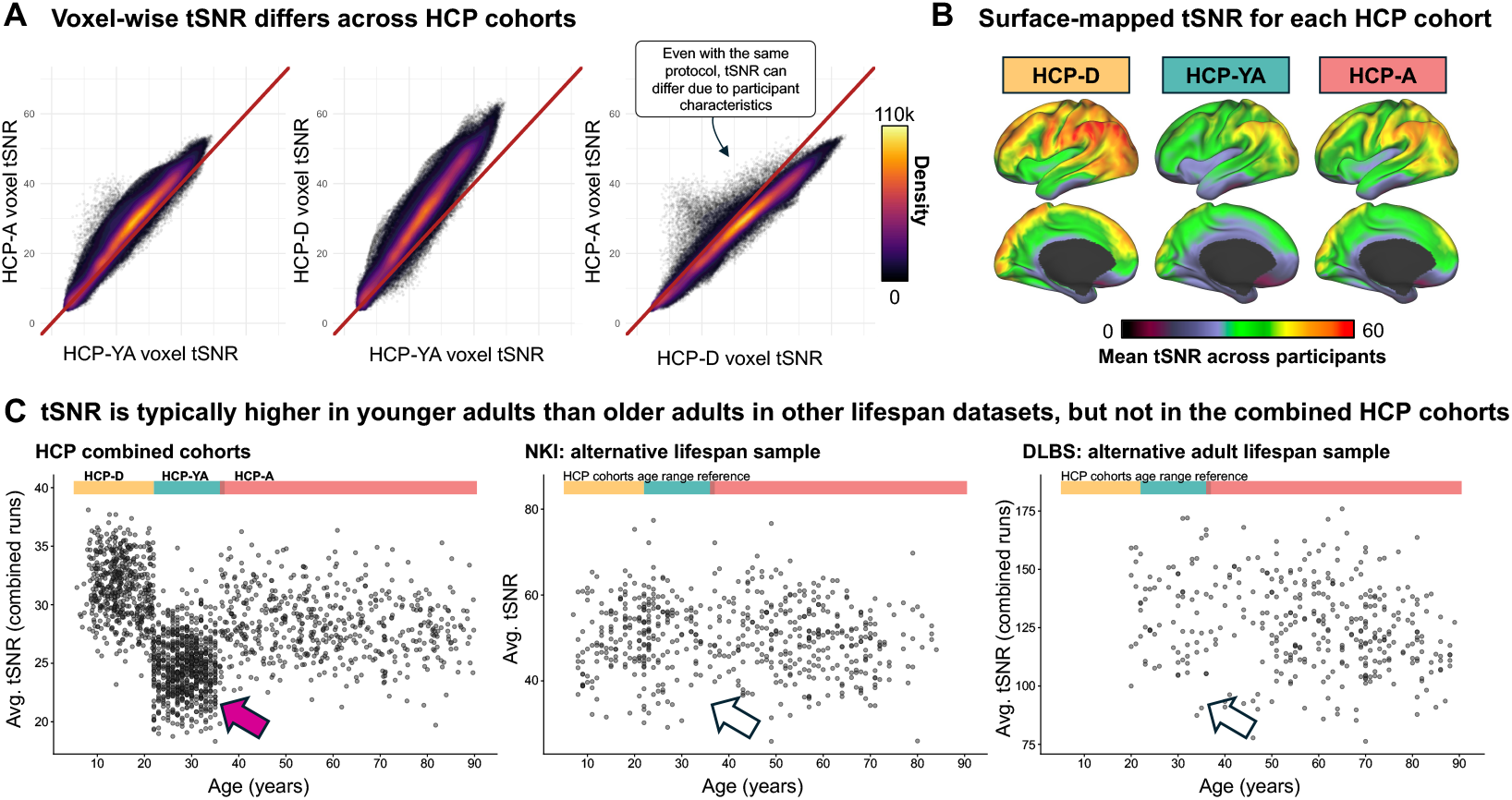
Temporal signal-to-noise ratio (tSNR) differs across cohorts. **(A)** Density scatter plots compare temporal signal-to-noise ratio (tSNR) of whole brain voxels (brain masked) between the three HCP cohorts. The red diagonal line is the identity (origin of 0, slope of 1). Deviation from the identity line indicates how tSNR differs across cohorts, where HCP-D/A exhibit higher tSNR compared to HCP-YA. All cohorts’ data were processed locally using HCP-pipeline v4.2 with the same software packages (FreeSurfer 6.0, FSL 6.0.4, MATLAB 2021a, Workbench 1.5.0). **(B**) Surface-mapped average tSNR values of each cohort. **(C)** Age by average resting-state tSNR (averaged across all voxels after brain mask) in three distinct datasets (HCP-combined cohorts, NKI, DLBS). (**C-left)** The HCP-YA cohort show a clear difference in tSNR compared to the HCP-D and HCP-A cohorts, suggesting a potential cohort-level confound in tSNR. Colored bars above the plot indicate HCP-cohort age range (note that HCP-YA and HCP-A have minor overlap in age). The magenta arrow highlights a marked reduction in tSNR across HCP-YA cohort. **(C-middle & right)** tSNR across age from two independent datasets with a lifespan sample (NKI; middle) and adult lifespan sample (DLBS; right) are depicted and show no marked reduction in tSNR among younger adults (see clear arrows). Colored bars in NKI and DLBS plots indicate HCP cohorts age ranges for reference.

#### 2.4.2 Measure of functional brain network organization

Each individual’s network organization was evaluated using three mean RSFC measures (mean RSFC across the whole brain, mean within-system RSFC, mean between-system RSFC) and four graph-theoretical network measures (brain system segregation, modularity, participation coefficient, and clustering coefficient).

For measures where system assignment (i.e., community) was required, predefined large-scale brain systems associated with each brain atlas were used. For both the surface-based datasets (HCP and DLBS) and the volumetric-based dataset (NKI), nodes were assigned to the Power et al. canonical functional networks (Chan et al., 2014; Power et al., 2011). These system assignments were used to compute within- and between-system functional connectivity, system segregation, and participation coefficient.

##### Brain system segregation

System segregation measures the degree to which the functional brain network is organized into distinct systems (Chan et al., 2014; Wig, 2017). Whole brain system segregation was calculated as the difference between mean within-system and mean between-system correlations as a proportion of mean within-system correlation, as noted in the following revised formula in (Chan et al., 2021):

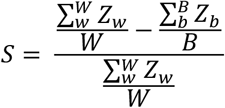

where *Z* is a Fisher’s *z*-transformed correlation value, representing an edge between a pair of nodes. *Z*_w_ denotes the edges (correlations) between node pairs belonging to the same system (*within-system correlations), Z*_b_ denotes the edges between node pairs belonging to different systems (*between-system correlations*), *W* is the total number of within-system edges across all subnetworks, and *B* is the total number of between-system edges across all subnetworks. A higher value of *S* indicates greater segregation of the brain into distinct functional systems.

For system segregation, negative correlations were excluded from the brain network matrix (i.e., negative values were set to zero), following previous reports (Chan et al., 2014, 2018). When mean within-system 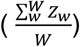 and mean between-system functional correlation 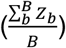were used separately as RSFC summary measures, positive and negative edges were retained.

##### Modularity

Modularity (Q) quantifies the extent to which a network can be partitioned into distinct modules or communities (Newman, 2004b). Weighted modularity is defined by the following formula (Newman, 2004a):

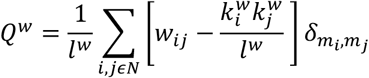

where *l*^*w*^ is the sum of all weighted edges in a network, *w*_*ij*_ is the weighted edge between nodes *i and j*, 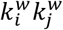are the total weighted degree (strength) of nodes *i and j, m*_*i*_ is the module containing node *i*, and *δ*_*mi,mj*_ = 1 if *m*_*i*_ = *m*_*j*_, and ‘0’ otherwise. Modularity was computed from 4% edge-density matrices.

##### Participation coefficient

The weighted participation coefficient (PC) quantifies to what extent a node interacts with nodes from other functional systems, proportional to its overall connection (total weighted degree) (Guimerà & Amaral, 2005). A node with higher PC indicates more between-system connections relative to the node’s total connectivity. The formula for calculating a node’s weighted participation coefficient 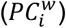 is:

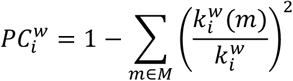

where 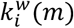 represents the sum of weighted connections from node *i* to nodes belonging to system *m* (excluding the system to which node *i* itself belongs), and 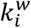 represents the total sum of weighted connections for node *i*. A higher PC for a given node indicates proportionally greater connectivity with nodes in other systems relative to its total connections. The PC of each node was computed from 4% edge-density matrices. The mean PC used in the current work was obtained by averaging 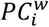across all nodes.

##### Clustering coefficient

The clustering coefficient (CC) quantifies the extent to which the neighbors of a node are also connected to one another, reflecting the presence of local clustering or “triangles” in the network (Watts & Strogatz, 1998). The weighted clustering coefficient quantifies the strength of triangle formation around nodes in weighted networks (Onnela et al., 2005).

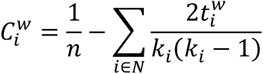

where 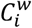 is the weighted clustering coefficient, *n* is the total number of nodes, 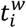denotes the total strength of the weighted triangles around node *i*, and *k*_*i*_ is the degree of node *i*. A higher 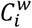 indicates stronger local interconnectedness when considering edge weights. The CC of each node was computed from 4% edge-density matrices. The mean CC used in the current work was obtained by averaging 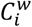 across all nodes.

### 2.5 Data harmonization

To integrate data across the three HCP cohorts (HCP-YA, HCP-D, and HCP-A), we applied CovBat (Chen et al., 2021, 2022) and ComBat-GAM (Pomponio et al., 2020) for harmonization. Rather than focusing solely on site-specific effects, we defined a batch variable to reflect protocol/scanner-model differences between HCP cohorts (HCP-YA: TR=720ms, LR/RL encoding, custom Skyra scanner vs. HCP-D/A: TR=800ms, AP/PA encoding, Prisma scanner). This grouping of participants by acquisition protocol/scanner-model was used as the batch factor in harmonization models, capturing the major acquisition differences between HCP-YA and the later datasets (see **SI 1.10** for differences in harmonization targeting different batch variables). In both approaches, the harmonization method removed batch-related variation while retaining variance associated with selected covariates. Based on their biological relevance, age and sex were included as selected covariates in the harmonization models and explicitly preserved to allow targeted examination of age-related trends.

#### 2.5.1 CovBat

CovBat (Chen et al., 2022) was used to adjust batch effects in RSFC matrix edges across participants. CovBat is an extension of ComBat, which aims to remove batch effects in mean and variance of features; CovBat also aligns the inter-feature covariance, which is useful for accurately comparing functional connectivity patterns across cohorts. Age and sex were included as covariates. Accordingly, RSFC matrices were harmonized such that variance associated with age and sex was preserved. CovBat was implemented in R using the *CovBat_Harmonization* package.

#### 2.5.2 ComBat-GAM

ComBat-GAM (Pomponio et al., 2020) was used to adjust batch effects at the level of RSFC and graph summary measures (e.g., mean RSFC, system segregation). ComBat-GAM is an extension of ComBat, but incorporates generalized additive models (GAMs) to account for nonlinear effects of covariates. Like ComBat, ComBat-GAM removes unwanted variance associated with batch or site effects while preserving variance related to covariates of interest (e.g., age, sex). However, whereas traditional ComBat assumes linear relationships between covariates and the data, ComBat-GAM uses smooth, nonparametric functions to flexibly model nonlinear trajectories across the lifespan or other continuous variables. ComBat-GAM was implemented in R using the *ComBatFamily* package.

### 2.6 Statistical analysis

Age-related trajectories of RSFC measures and graph network metrics were modeled using generalized additive models (GAMs) implemented in the *mgcv* package in R (Wood, 2017). GAMs extend generalized linear models by allowing the inclusion of smooth, non-linear functions of predictors, enabling flexible estimation of age trends without assuming a specific parametric form. For each measure of interest (e.g., mean RSFC, within-system RSFC, system segregation), a GAM was fitted with age as a continuous predictor modeled with thin-plate regression splines. In the present analysis, the goal of the GAM fits was to provide a smooth visualization of irregular age-related trends in RSFC measures, rather than to serve as a formal statistical test or model of developmental or aging effects.

### 2.7 Software

Statistical analysis and data visualization were conducted in R 4.4.1 using the following packages: *ComBatFamily* (0.2.1), *ggplot2* (3.5.1), *ggpointdensity* (0.1.0), *mgcv* (1.9), and *stats* (4.4.1). Visualizations of tSNR on the cortical surfaces were generated using Connectome Workbench (1.5.0).

## 3. RESULTS

### 3.1 tSNR differs between HCP cohorts

In time series data such as those collected with fMRI, temporal signal-to-noise ratio (tSNR) reflects the stability of the BOLD signal over time, and is commonly used to evaluate signal quality (Jamil et al., 2021; Krüger & Glover, 2001; Parrish et al., 2000; Welvaert & Rosseel, 2013). To minimize the influence of downstream data preprocessing steps on tSNR, the same version of the HCP-pipeline fMRI volumetric pipeline (i.e., distortion correction, realignment, MNI registration, intensity normalization, brain masking) was used to preprocess all three HCP cohorts. The tSNR of processed time series prior to motion-censoring and surface-mapping was compared (see **SI 1.1, Fig. S1** for a comparison of tSNR extracted from the “Preprocessed Recommended fMRI Data” provided by the HCP team for each cohort). For each cohort, only participants with minimal head motion (mean filtered framewise displacement [fFD] < 0.04) and whose average tSNR–across all runs–did not fall below 3 SDs of their cohort’s mean were included to reduce potential confounding effects of motion and extreme tSNR cases on fMRI signal quality (see **SI 1.2, Fig. S2A** for high-motion participants that were removed from the main analysis). Within each cohort, tSNR was calculated for each run of MNI-registered resting-state fMRI data, computed voxel-wise across time, and then averaged across all participants.

The cohort-average tSNR maps were compared voxel-wise in **Fig. 1A**. Both HCP-A and HCP-D had higher tSNR than HCP-YA: HCP-A vs. HCP-YA (tSNR_mean_=27.16 vs. 23.09, *t*=134.48, *p*<.001), HCP-D vs. HCP-YA (tSNR_mean_=29.53 vs. 23.09, *t*=193.04, *p*<.001). Because HCP-YA included a larger sample, additional comparisons were made by subsampling all three cohorts to compare a narrower age range with approximately matched sample sizes, which showed similar cohort-level differences (see **SI 1.3, Fig. S3**). **Fig. 1B** depicts the cohort tSNR mapped to the 32k fs_LR surface, in which lower tSNR values were also evident across multiple regions in the HCP-YA cohort, compared to the other two cohorts.

Although these tSNR differences may appear subtle and could be within the range observed across participants collected under identical protocols (as indicated by the right panel of **Fig. 1A** comparing HCP-D to HCP-A), a clearer cohort confound emerged when examining tSNR as a function of cohort and age. When examining average tSNR (combined across functional runs) as a function of participant age **(Fig. 1C-left)**, the HCP-YA cohort showed a clear reduction in tSNR compared to HCP-D and HCP-A.

It should be noted that although HCP-D and HCP-A shared the same scanning protocol, the comparison between the two showed that HCP-D, which includes children, adolescents, and younger adults, exhibited higher tSNR than HCP-A. This suggested potential age-related decreases in tSNR (tSNR_mean_=29.53 vs. 27.16, *t*=68.56, *p*<.001; also see **SI 1.4, Fig. S4** for age-related differences in tSNR across adulthood in HCP cohorts and independent datasets, and **SI 1.5, Fig. S5** for distribution comparisons of tSNR across HCP cohorts). Overall though, while some signal quality reduction occurs with age in adulthood (Huettel et al. 2001), it was unexpected that the HCP cohort with the oldest adults (HCP-A) exhibited higher tSNR than younger adults in HCP-YA.

To ensure that the transition from the developmental cohort to the aging cohort (i.e., young adulthood) was not inherently a segment of the lifespan associated with lower tSNR, two independent resting-state fMRI datasets with a wide age range were analyzed (**Fig. 1C-middle**: Nathan Kline Institute Rockland Sample [NKI] and **Fig. 1C-right**: Dallas Lifespan Brain Study [DLBS]). Younger adults in both datasets showed no marked reduction in tSNR compared to younger and/or older participants.

Overall, HCP-YA exhibited significantly lower tSNR across the entire sample when compared to both the developmental and aging cohorts (HCP-D and HCP-A). Younger adults in other datasets did not exhibit reduced tSNR compared to their developmental/aging counterpart, indicating that reduction in tSNR observed in HCP-YA was unlikely to be due to a true age-related effect, but rather, cohort-related differences due to variations in MRI acquisition across HCP datasets. The observed cohort-related effect was a type of “batch differences” (Hu et al., 2023), representing non-biological variability introduced by differences in acquisition protocol and scanner model. Not only can these batch differences shift the distribution and scaling of tSNR, but they could also impact downstream RSFC-derived metrics, potentially introducing atypical age-related trajectories unless appropriately harmonized.

### 3.2 Impact of batch differences on RSFC network measures across HCP cohorts

We evaluated the impact of batch differences on age-related differences in measures of RSFC network organization across HCP cohorts (**Fig. 2A**). We also examined these relationships in the NKI (**Fig. 2B**) and DLBS (**Fig. 2C**) datasets for reference. We examined general measures of RSFC mean correlations (whole brain, within-system, and between-system correlation) along with several commonly used network measures that summarize RSFC network topology both at the whole-brain level and at the node level. For visualization, generalized additive models (GAMs) were fit to individual observations to provide smooth age-related trajectories; these curves were intended to highlight irregularities in age patterns.

**Figure 2.**
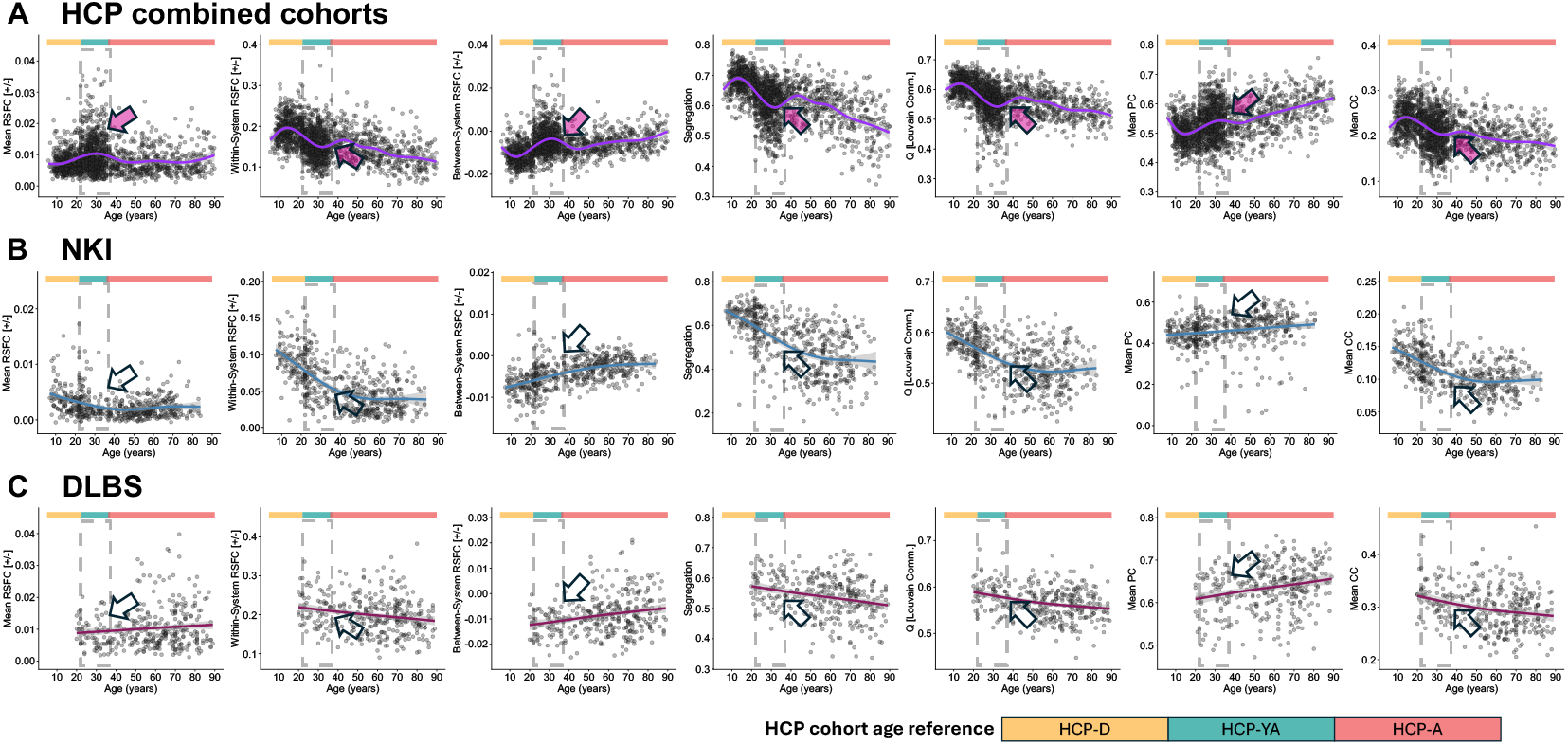
Resting-state functional correlation (RSFC) structure and derived graph measures show notable differences in the HCP-YA compared to the other two cohorts. These differences are not observed within the same young-adult age range in the NKI (lifespan) and DLBS (adult lifespan) datasets. Generalized additive models (GAMs) were fitted to visualize age-related trends for seven measures of large-scale functional network organization (left to right): mean RSFC, within-system RSFC, between-system RSFC, system segregation, modularity (Q), mean participation coefficient (PC), and mean clustering coefficient (CC). Three distinct data collection efforts which span large segments of the lifespan are visualized: HCP, NKI, and DLBS. Colored bars above graphs indicate the different HCP-cohorts (yellow=HCP-D, teal=HCP-YA, salmon=HCP-A) and dashed gray lines demarcate the 25-37 HCP-YA age-range for easy visualization of distinct properties of the HCP-YA data. The colored bars and grey lines are maintained in the NKI and DLBS plots for easy reference. **(A)** HCP cohorts show strong cohort-related deviations in these RSFC measures (see pink arrows). For example, the HCP-YA cohort exhibits inflated mean and between-system RSFC alongside reduced within-system RSFC and system segregation, leading to atypical age-related trajectories. These deviations in young-adult functional connectivity are not evident in either the **(B)** NKI lifespan or **(C)** DLBS adult-lifespan dataset, each of which shows smoother, biologically plausible age-related trends, with younger adults exhibiting higher segregation, modularity, and clustering coefficient than older adults (see hollow arrows).

Focusing first on the HCP cohorts, the mean RSFC correlations were higher in HCP-YA than the participants in a similar age range from the other two HCP cohorts (i.e., the upper age range in HCP-D and the lower age range in HCP-A). As most edges in an RSFC matrix represent correlations between functional systems, the increase in whole-brain RSFC in HCP-YA was largely driven by elevated between-system RSFC (see **Fig. 2A, 3**^**rd**^ **panel**). Conversely, within-system RSFC correlation was lower in HCP-YA compared to the other two HCP cohorts. This pattern—elevated between-system and overall RSFC alongside reduced within-system RSFC—was not observed in a comparable young-adult age range in the two reference datasets (compare pink vs. hollow arrows and dashed rectangular boxes highlighting the young-adult range in **Fig. 2A-C**). To statistically evaluate the differences across cohorts, the oldest HCP-D participants (aged 20–21y; n=77) were compared to the youngest HCP-YA participants (aged 22–23y; n=85), and the youngest HCP-A participants (aged 36–38y; n=29) were compared to the oldest HCP-YA participants (aged 35–37y; n=45). Across the seven graph measures, the 22–23 years old from HCP-YA were statistically different than the 20–21 years old from HCP-D (see 1^st^ row of **Table 1;** also see **Table S1 [SI 1.6]** for confidence intervals of the mean estimates). This between-group difference was also observed in 4 out of the 7 measures among middle-aged adults in HCP-YA vs. HCP-A (see 2^nd^ row of **Table 1** and **Table S1**). Notably, similar age-accompanied differences were not observed in the NKI or DLBS datasets (row 3-6 of **Table 1**).

**Table 1.** Comparing RSFC measures between adjacent age intervals in HCP cohorts, independent datasets (NKI, DLBS), and harmonized HCP cohorts (CovBat and ComBat-GAM)

|  | Mean RSFC | Mean within-system RSFC | Mean between-system RSFC | System Segregation | Modularity (Q) | Mean Participation Coefficient | Mean Clustering Coefficient |
| --- | --- | --- | --- | --- | --- | --- | --- |
| **HCP cohorts before data harmonization** |  |  |  |  |  |  |  |
| HCP-D (20-21y \| n=77) vs. HCP-YA (22-23y \| n=85) | *** | *** | *** | *** | *** | * | ** |
| HCP-YA (35-37y \| n=45) vs. HCP-A (36-38y \| n=29) | * | ns | *** | *** | *** | ns | ns |
| **Independent datasets** |  |  |  |  |  |  |  |
| NKI (20-21y \| n= 49) vs. NKI (22-23y \| n= 22) | ns | ns | ns | ns | ns | ns | ns |
| NKI (33-36y \| n= 26) vs. NKI (37-40y \| n = 22) | ns | ns | ns | ns | ns | ns | ns |
| DLBS (20-21y \| n = 12) vs. DLBS (22-25y \| n = 13) | ns | ns | ns | ns | + | ns | ns |
| DLBS (33-36y \| n = 20) vs. DLBS (37-45y \| n = 23) | ns | ns | ns | ns | ns | ns | ns |
| **HCP cohorts after data harmonization** |  |  |  |  |  |  |  |
| <u>CovBat</u><br>HCP-D (20-21y \| n=77) vs. HCP-YA (22-23y \| n=85) | ns | ns | ns | ns | * | ns | ns |
| <u>CovBat</u><br>HCP-YA (35-37y \| n=45) vs. HCP-A (36-38y \| n=29) | ns | ns | ns | ns | * | ns | ns |
| <u>ComBat-GAM</u><br>HCP-D (20-21y \| n=77) vs. HCP-YA (22-23y \| n=85) | ns | ns | ns | ns | ns | ns | ns |
| <u>ComBat-GAM</u><br>HCP-YA (35-37y \| n=45) vs. HCP-A (36-38y \| n=29) | ns | ns | ns | ns | ns | ns | + |

The age ranges for the comparison datasets (NKI and DLBS) were extended in certain intervals to increase participant coverage. This expansion is expected to increase, rather than reduce sensitivity to age-group effects.
+p<.10, \*p < .05, \*\* p < .01, \*\*\* p < .001
See **Table S1** for confidence intervals of RSFC measures' mean estimates

Collectively, these observations indicated that the atypical RSFC patterns in HCP-YA were unlikely to reflect true neurobiological differences. Instead, the differences across HCP cohorts were consistent with a batch effect arising from methodological differences in protocol/scanner-model (i.e., HCP-YA being collected on a different scanner and with different scan parameters compared to HCP-D and HCP-A; also see **SI 1.7, Fig. S6** showing how tSNR and RSFC network measures varied across three resting-state acquisitions from NKI). We next examined whether and how these differences resulted in biased measures of RSFC network topology.

The last four panels of **Fig. 2A-C** illustrate four graph measures of brain network organization. Two are whole-brain/network level measures: system segregation (Chan et al., 2014) and modularity determined by Louvain community detection (Newman, 2004b). Two are summary node-level network measures: mean participation coefficient (Guimerà & Amaral, 2005) and mean clustering coefficient (Watts & Strogatz, 1998). In the NKI and DLBS datasets (see rows 3-6 of **Table 1** and **Table S1**), younger adults showed higher system segregation, modularity, and clustering coefficient than adults who were slightly older (even as early as early middle-age), whereas younger adults from HCP-YA showed the opposite pattern when compared to middle-aged adults in HCP-A. Mean participation coefficient exhibited the reverse direction of effects, where younger adults had lower participation coefficient than early middle-age adults in NKI/DLBS, but the opposite effect was observed across HCP cohorts.

### 3.3 Cohort differences in RSFC can be mitigated with data harmonization

The observed batch effect (Hu et al., 2023) in RSFC network measures can be corrected with various methods. The targeted ‘batch’ effect to be adjusted for is protocol/scanner-model differences. We included adjustments at the feature level (i.e., region-to-region correlations) that make up the correlation matrix (e.g., CovBat; Chen et al., 2021), and at the summary measure level (i.e., graph measures of brain network topology; ComBat-GAM; Pomponio et al., 2020). Previous studies have applied different data harmonization approaches to correct for batch differences (e.g., ComBAT, Yu et al., 2018; CovBAT, Chen et al., 2022; SMA, Y.-W. Wang et al., 2023), or modeled batch-differences directly in the statistical analyses (e.g., Bethlehem et al., 2022). The present work did not aim to exhaustively compare methods for batch adjustment (see Wang et al. 2023), but rather to (1) illustrate how uncorrected batch-related differences can influence downstream RSFC measures, and (2) demonstrate how harmonization methods such as CovBat or ComBat-GAM can help alleviate these effects.

We compared the same summary RSFC measures that were depicted in **Fig. 2**, but harmonized the three HCP datasets with two separate methods: (1) CovBat corrected batch effects at the RSFC edge level, in which the summary graph measures were calculated with the adjusted RSFC matrices; (2) ComBat-GAM corrected batch effects at the summary feature level. Both methods were applied to target the specific batch differences between protocol/scanner-model (HCP-YA vs. HCP-D/A), and preserved the following covariates: age and sex—this ensured that meaningful variance with age was retained while removing unwanted batch effects.

Following harmonization using CovBat or ComBat-GAM, the HCP-YA data exhibited reduced deviations from the established age-related trends observed in NKI and DLBS (**Fig. 3** vs. **Fig. 2B & 2C**; also see **SI 1.8** for similar comparisons using alternate processing considerations). CovBat effectively adjusted the age trajectories of mean RSFC, within- and between-system RSFC, system segregation, mean participation coefficient, and mean clustering coefficient, with moderate improvement for modularity (see rows 7-8 of **Table 1** and **Table S1** for comparisons of adjacent age-intervals across HCP cohorts after CovBat). This resulted in a closer alignment of HCP-YA data with the expected age-trend, with fewer abrupt transitions in the age-curve between HCP-YA and other cohorts. Similarly, ComBat-GAM reduced the deviations in HCP-YA across all measures (see rows 9-10 of **Table 1** and **Table S1**). Supplemental analyses showed that alternative harmonization batching strategies were effective when the batching structure separated HCP-YA from HCP-D/A, thereby preserving the key protocol/scanner-model distinctions. In contrast, batching strategies that combined HCP-YA participants with HCP-D/A participants within the same batch were not effective, even when those batches reflected other potentially relevant factors but failed to batch the protocol/scanner-model differences (e.g., geographic site; see **SI 1.9 and SI 1.10**).

**Figure 3.**
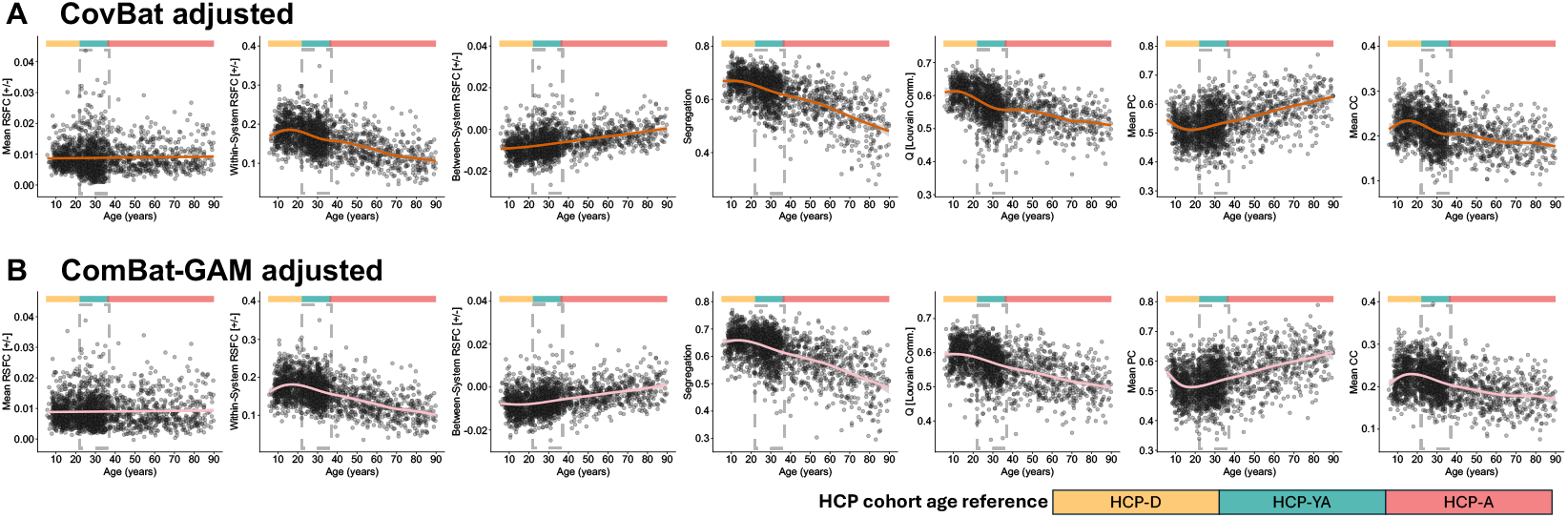
Data harmonization focused on sources of batch differences improves measurement of RSFC and network measures across the lifespan. Scatterplots depicts GAM fit of adjusted summary RSFC or network measures across age following batch adjustment (left to right): mean RSFC, within-system RSFC, between-system RSFC, system segregation, modularity (Q), mean participation coefficient (PC), and mean clustering coefficient (CC). **(A)** CovBat harmonized the mean, variance, and covariance of the RSFC matrix edges across datasets. **(B)** ComBat-GAM harmonized the summary features, adjusting for mean and variance. The grey box in each panel marks the HCP-YA age range (as in **Figure 2**) to facilitate comparison to unadjusted curves.

## 4. DISCUSSION

### 4.1 General discussion

If taken at face value, measures of resting-state network organization calculated from the unharmonized data from the three HCP cohorts imply an unlikely developmental and aging trajectory where younger adults exhibit reduced network segregation and modularity relative to both adolescent and midlife individuals. This pattern is inconsistent with previously published work (Betzel et al., 2014; Chan et al., 2014; Deery et al., 2023; Geerligs et al., 2015; Pedersen et al., 2021), and with the independent lifespan cohorts examined here. These discrepancies indicate that, without accounting for protocol-related batch differences, analyses combining HCP cohorts can yield qualitatively misleading–and even inverted–interpretations of developmental and aging trajectories. More broadly, this issue is not unique to HCP, but applies to any multi-cohort lifespan study incorporating protocol or hardware differences or changes over time, where unmodeled signal differences can distort apparent developmental or aging trajectories.

Although prior work has applied harmonization approaches to the HCP datasets (e.g., Y. Wang et al., 2025), the effects of these procedures and the consequences of failing to apply them have not been directly demonstrated. Our results provide a direct evaluation of how cohort differences manifest in network measures and show that harmonization materially impacts the characterization of lifespan trajectories of brain network organization.

The differences observed in HCP-YA relative to HCP-D and HCP-A illustrate how batch effects that are observable in signal characteristics, particularly tSNR, can systematically bias functional connectivity estimates and network measures. The batch effect being targeted in the present work was focused on the combined effect of HCP-YA having a different protocol (scan acquisition parameters) and scanner model (hardware) compared to HCP-D and HCP-A (see **SI 1.9-1.10, Fig. S9** for analyses using geographical ‘site’ or a combination of ‘site + cohort’ and ‘site + scanner-model’, showing that harmonization strategies that differentiated HCP-YA into its own batch yielded acceptable results). With proper harmonization, researchers can avoid misinterpreting methodological variation as genuine neurobiological differences.

We have focused on data collected under several HCP initiatives that support lifespan studies of brain and cognition; these findings do not diminish the value of the HCP datasets, which remain exceptional in both quantity and quality of data. Rather, by incorporating harmonization approaches and demonstrating that batch effects are adequately addressed, researchers can more confidently leverage this powerful set of studies and interpret the observed developmental and aging trajectories.

The present work demonstrated that HCP-YA exhibited substantially lower tSNR, and this reduction in tSNR systematically biased RSFC estimates and derived network measures. This resulted in RSFC measures obtained from HCP-YA, when examined in conjunction with HCP-D and HCP-A, diverging from expected developmental and aging trajectories observed in two independent datasets (NKI, DLBS). Specifically, HCP-YA data showed higher between-system and overall RSFC but lower within-system RSFC, along with inverted trajectories for network-level measures such as system segregation, modularity, participation coefficient, and clustering coefficient. These differences are unlikely to reflect genuine neurobiological effects, as they were not observed in other lifespan cohorts; notably, similar differences were still evident across HCP cohorts when overlapping ages were examined (see comparisons in **Table 1** and **Table S1**). Instead, the differences across HCP cohorts highlight a cohort-specific batch effect driven, in part, by scanner hardware and acquisition protocols.

These results underscore how methodological variability, if not adequately accounted for, could be misinterpreted as developmental or aging effects. Without data harmonization, investigators could mistakenly interpret the atypical HCP-YA patterns as altered measures of brain function during young adulthood, especially if comparisons are constrained to adjacent segments of the lifespan (e.g., HCP-D and HCP-YA or HCP-YA and HCP-A), which would obscure the u-shaped or inverted u-shaped patterns described here. More broadly, this type of concern is not limited to RSFC measures, but can also extend to other BOLD-derived metrics, including task-evoked activations, BOLD fluctuation-amplitude metrics, and cerebrovascular reactivity, which have been shown to be sensitive to differences in hardware, acquisition, or preprocessing methods (Brown et al., 2011; Friedman et al., 2008; Liu et al., 2021; X. Wang et al., 2021). The NKI and DLBS datasets both showed a more continuous pattern of lifespan differences in RSFC network organization, without the pronounced young-adult deviation observed in unharmonized HCP data. These findings reinforce the conclusion that the HCP-YA deviations reflected a cohort-specific batch effect rather than a genuine biological phenomenon.

The acquisitions and processing pipelines for the two reference datasets (NKI and DLBS) differed in several respects from those used in HCP cohorts. DLBS featured a longer TR (2000ms) and 3mm isotropic voxels, resembling many legacy fMRI datasets; in terms of preprocessing, we employed the same surface-based node set as what was used in HCP. The short-TR acquisition from NKI permitted an alternative comparison lifespan sample with different data acquisition parameters (see **SI 1.7** for other NKI acquisitions), and we used a volumetric node set along with several additional minor differences in denoising strategies (e.g., aCompCor with 5 white matter components instead of a single white matter mask during nuisance regression; major processing choices were shared with other datasets used in this work: global signal regression, high-motion volume censoring). The inclusion of these variations in node definitions and processing steps demonstrates the consistency of age-related RSFC trajectories across these independent participant cohorts and data acquisition parameters, supporting their utility as reference cohorts for evaluating the protocol/scanner-model effects observed in HCP-combined cohorts and impacts of harmonization.

These results have broader implications for lifespan studies that rely on integrating multiple cohorts or datasets. Combining multiple datasets can increase statistical power and expand generalizability through increased sample size and sample diversity, but it also introduces heterogeneity that can bias inferences about developmental and aging trajectories. Methodological variation can also be introduced in single cohort studies through hardware/software upgrades or changes in study procedures (e.g., procedural alterations to improve participant retention/comfort). Researchers should carefully account for differences in hardware, sequence design, and acquisition quality when combining datasets. Systematic data-quality assessment that explicitly checks for cross-cohort differences should be part of standardized data processing workflows.

In this work, we primarily focused on mitigating the effect of protocol/scanner-model. However, harmonization approaches could in principle target factors such as cohort (e.g., HCP-D, HCP-YA, HCP-A), site (e.g., geographical locations), or a combination of multiple factors (e.g., sites within a cohort). Supplemental results in **SI 1.10** include analyses showing that harmonization strategies targeting different factors were similarly effective provided that protocol/scanner-model differences were explicitly modeled in the batching structure. For example, harmonization based on geographic site alone resulted in batching together data acquired under different protocols and scanner models, because some geographical sites collected data for all three HCP cohorts. As a result, protocol/scanner-related differences were not properly isolated, and the atypical lifespan trajectory remained evident even after harmonization.

Signal quality could be measured in various ways; here, we focused on tSNR as it captures the aggregated impact of multiple acquisition parameters on signal magnitude and variance. While tSNR is mathematically related to BOLD variability (i.e., BOLD variability is the denominator of the tSNR formula), which has been linked with biologically meaningful neural signal (e.g., Grady & Garrett, 2018; also see review, Waschke et al., 2021), substantial work has shown that the inclusion of mean signal makes tSNR sensitive to acquisition-related factors that impact signal quality and noise characteristics (Jamil et al., 2021; Krüger & Glover, 2001; Triantafyllou et al., 2005; Welvaert & Rosseel, 2013). Furthermore, the observed pattern of tSNR differences across HCP cohorts did not follow a plausible lifespan trajectory and was inconsistent with age-related patterns observed in two independent datasets, suggesting that the observed differences were primarily driven by acquisition-related factors rather than age-related neurobiological differences. However, we also acknowledge that poor tSNR itself is not always an artifact and could represent true biological differences (e.g., reduced cerebral blood flow and vascular reactivity with age affect BOLD signal; Wright & Wise, 2018; Zimmerman et al., 2021). Therefore, we did not use any data correction approaches that directly used tSNR as a feature or covariate (e.g., controlling for tSNR as a covariate in a linear model). Future work should continue to evaluate other complementary signal-quality metrics and harmonization strategies that can separate biologically meaningful variance in BOLD signal from acquisition-related biases.

Although we did not exhaustively compare harmonization strategies, the current findings underscore the necessity of applying data harmonization for valid cross-dataset analyses in lifespan research. CovBat and ComBat-GAM were used here as illustrative examples, the best approach may vary across datasets and outcome measures, as reflected by the differing effects of these methods across the RSFC network measures examined. In addition, the optimal stage for data harmonization during analysis (e.g., at the level of RSFC edges vs. summary network features) will depend on the specific research question and analytic goals. For example, CovBat was applied to RSFC matrices, in which all downstream network measures in **Fig. 3** were derived from the same harmonized connectivity estimates. Conversely, ComBat-GAM was applied separately to each summary measure. These approaches therefore differed not only in the harmonization method used, but also in the level of the analysis pipeline at which harmonization was performed. Researchers should therefore carefully consider both the harmonization approach and the stage of the analysis pipeline at which harmonization is performed, based on their specific datasets, outcome measures, and analytic goals.

### 4.2 Conclusion

The present work demonstrates that acquisition- and scanner-related differences across HCP cohorts introduce batch effects that systematically bias RSFC estimates and network measures. Without data harmonization, these artifacts can be misinterpreted as genuine developmental or aging differences. Comparisons with independent lifespan datasets (NKI, DLBS) support the conclusion that the atypical patterns in HCP-YA reflect methodological rather than biological effects. Importantly, we show that data harmonization can substantially mitigate these confounds and recover age-related trajectories more consistent with prior literature and independent cohorts. These findings highlight that while combining cohorts into larger datasets is valuable, both planned multi-cohort studies and secondary analyses that integrate multiple existing datasets necessitate rigorous quality assessment, preprocessing, and data harmonization to draw valid inferences about brain development and aging. In sum, data harmonization should be viewed not as a technical refinement, but as a prerequisite for valid cross-cohort inference in lifespan network neuroscience.

## DATA AND CODE AVAILABILITY

HCP-YA Data S1200 release data (downloaded March 2023) are available via the Human Connectome Project website (https://www.humanconnectome.org). HCP-D and HCP-A release 2.0 data (downloaded August 2022) are available via the NIMH Data Archive (NDA; https://nda.nih.gov). NKI processed data were downloaded from the NKI-RS repository (S3 bucket). DLBS data are available on OpenNeuro (https://openneuro.org/datasets/ds004856). CovBat was applied using R code available on https://github.com/andy1764/CovBat_Harmonization. RSFC matrix extraction and brain system segregation calculation were completed using code available on https://github.com/mychan24/system-segregation-and-graph-tools. Modularity, participation coefficient and clustering coefficient were calculated using the Brain Connectivity Toolbox (BCT; https://sites.google.com/site/bctnet).

## AUTHOR CONTRIBUTIONS

M.Y.C. designed the study; M.Y.C. analyzed the data; M.Y.C. and L.H., processed and quality-checked the data; G.S.W. supervised the research and analysis; M.Y.C. and G.S.W. wrote the manuscript; L.H. provided comments and edits on the manuscript.

## ACKNOWLEDGEMENT

The authors acknowledge the Texas Advanced Computing Center (TACC) at The University of Texas at Austin for providing computational resources that contributed to the research results reported in this paper.

URL: http://www.tacc.utexas.edu

