## Supplemental Information for "Systematic fMRI signal differences across cohorts alter lifespan trajectories of functional brain networks"

### 1. SUPPLEMENTARY RESULTS

#### *1.1 tSNR differences in HCP cohorts using "Preprocessed Recommended fMRI Data"*

Using "Preprocessed Recommended fMRI Data" provided by the HCP team, which included the HCP-pipeline fMRI volume processing with ICA FIX, and the HCP-pipeline fMRI surface processing), cross-cohort comparisons of tSNR were examined. Note that linear detrend (hp0) instead of effective linear detrend (hp2000) was used for HCP-A/D data due to increased computational efficiency.

Because the downloaded data included surface-mapped cortical and volumetric subcortical data, the tSNR of the surface vertices and subcortical voxels were each plotted independently in **Fig. S1**, allowing comparisons of the tSNR of processed data on surface vertices and subcortical voxels separately. HCP-YA still exhibited the lowest tSNR across the three cohorts.

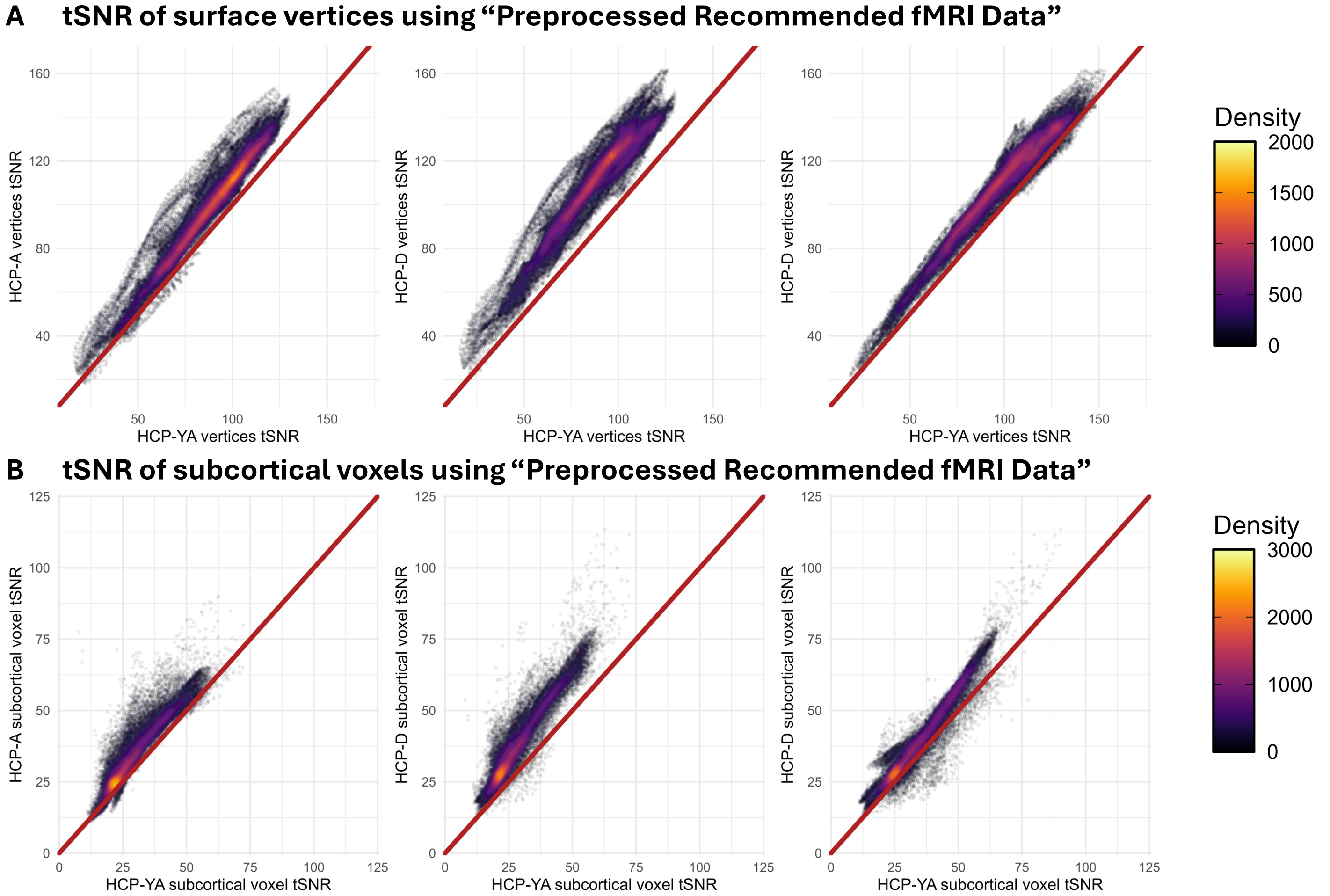

**Figure S1 – tSNR comparison using readily downloadable "Preprocessed Recommended fMRI Data" provided by the HCP.**
Across **(A)** surface vertices and **(B)** subcortical voxels, HCP-YA exhibits lower tSNR compared to HCP-D and HCP-A. With identical protocol and scanner model, HCP-D exhibits greater tSNR than HCP-A. Qualitatively, cross-cohort tSNR differences are similar with the locally-processed data used in the main analysis.

##### *1.2 High motion participants/volumes removed from tSNR analyses*

High motion was associated with weaker tSNR across all three HCP cohorts (below vs. above mean fFD of 0.04: HCP-D $t$=20.247, $p$ < .001; HCP-YA $t$ = 13.32, $p$ < .001; HCP-A $t$ = 13.32, $p$ < .001; see **Fig. S2A** with high-motion participants in red showing lower tSNR). To reduce the influence of motion on the interpretation of the results, high-motion participants (mean fFD > 0.04) were removed from the main analyses.

Here, tSNR was also recalculated with high-motion volumes of each of the retained participants removed from their preprocessed data. The cohort differences in tSNR between HCP-YA and HCP-D/A remained (HCP-YA vs. HCP-D: $t$ = 51.76, $p$ < .001; HCP-YA vs. HCP-A: $t$ = 28.33, $p$ < .001)

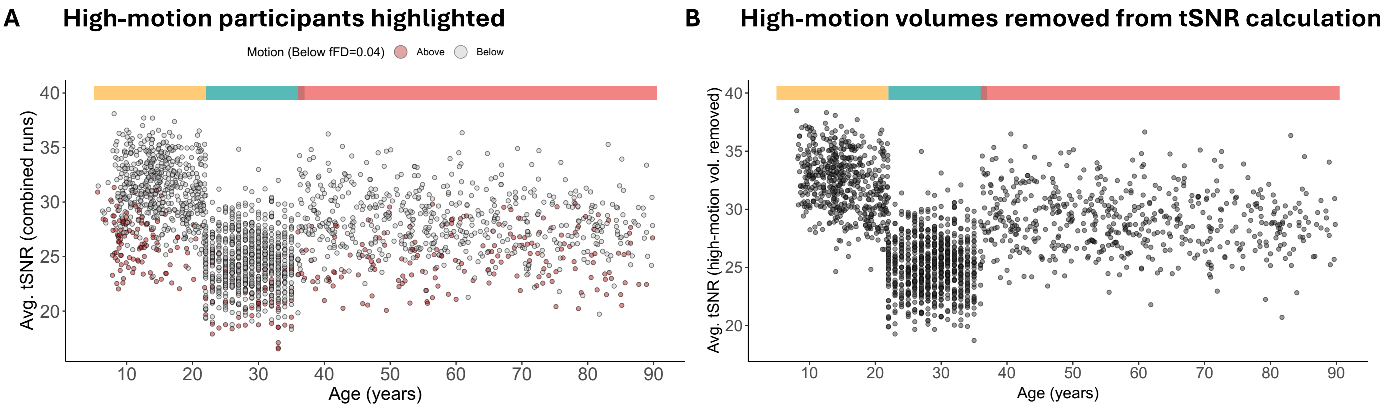

**Figure S2 – High motion participants from all HCP cohorts were removed from the main analyses.** **(A)** High motion participants (mean filtered FD> 0.04) are colored red, showing that they largely exhibit lower tSNR compared to participants without high motion. **(B)** When high-motion volumes were removed from the preprocessed data in participants who were retained for the main analyses, there remained a cohort difference in tSNR between HCP-YA and HCP-D/A.

##### *1.3 tSNR differences in HCP cohorts, restricting to participants in adjacent age-bins*

To reduce potential age-related effects in the tSNR comparison in the main text's **Fig. 1A**, here, we repeated the same analysis but only included participants most similar in age across cohorts (the same adjacent age intervals used in **Table 1**). In the HCP-D vs. HCP-YA comparison, HCP-D participants from age 20-21.5y and HCP-YA participants from 22-23y were included. In the HCP-YA vs. HCP-A comparison, HCP-YA participants from age 35-37y and HCP-A participants from age 36-38y were included. For HCP-D vs. HCP-A, the oldest HCP-D (20-21.5y) and youngest HCP-A (36-38y) participants were included (the smallest age gap between the two cohorts was ~15 years).

Similar to results including all participants in the main text, **Fig. S3** showed that the HCP-YA cohort exhibited a clear reduction in tSNR compared to HCP-D and HCP-A (i.e., deviation from the red diagonal line toward the side of HCP-D or HCP-A).

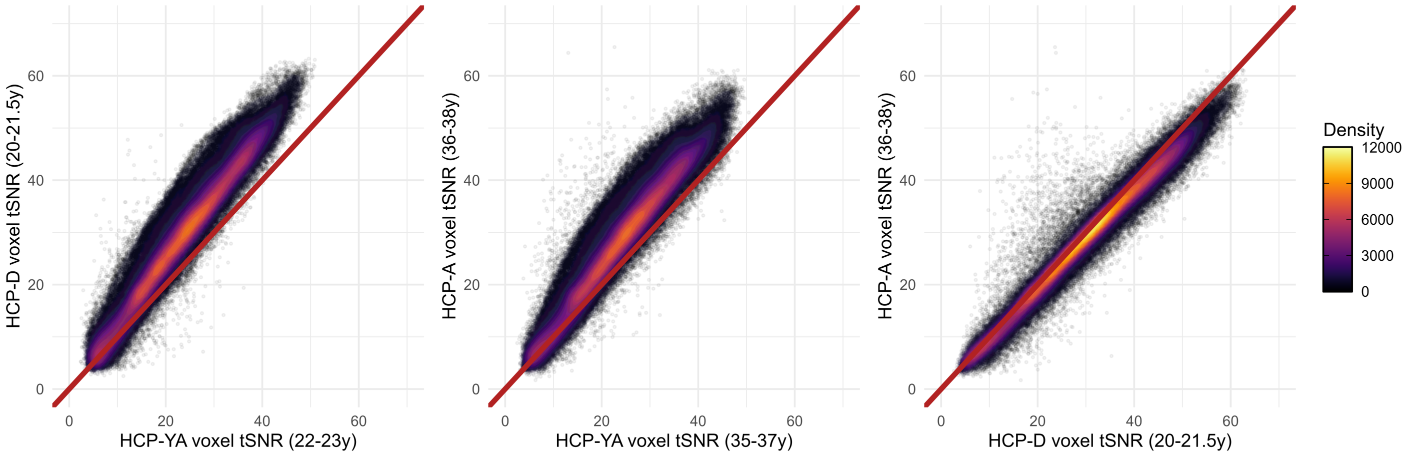

**Figure S3 – Approximately age-matched comparison still shows tSNR differences across cohorts.** Density scatter plots of temporal signal-to-noise ratio (tSNR) of whole brain voxels (brain masked) between the three cohorts are plotted. The red diagonal line is the identity (origin of 0, slope of 1). Deviation from the identity line indicates how tSNR differs across cohorts. HCP-D/A exhibit higher tSNR when compared to HCP-YA. To reduce potential age-related effects, only participants with closely matched ages across cohorts were included in each pairwise comparison. All cohorts' data were processed locally using HCP-pipeline v4.2 with the same software packages (FreeSurfer 6.0, FSL 6.0.4, MATLAB 2021a, Workbench 1.5.0).

#### *1.4 tSNR reduces with age in adult cohorts that share the same scan protocols*

Age-related differences in tSNR were examined in HCP cohorts and revealed a significant age-related decline in tSNR both in HCP-D (*r* = -0.16, *p* < .001) and HCP-A (*r* = -0.13, *p* = .002), but not in HCP-YA (*r* = 0.002, *p* = .962). To examine whether this relationship was evident in multiple adult cohorts, a subset of participants from the NKI lifespan dataset aged 18y or older and the DLBS adult lifespan dataset were examined. These comparisons revealed a significant decline in tSNR across the adult lifespan in these additional datasets: NKI adult subset: *r* = -0.14, *p* < .001; DLBS: *r* = -0.21, *p* < .001. These results demonstrate that the gradual decline in tSNR with age in adulthood was consistently observed across datasets, in contrast to the cohort-specific shifts observed between HCP-YA and HCP-A.

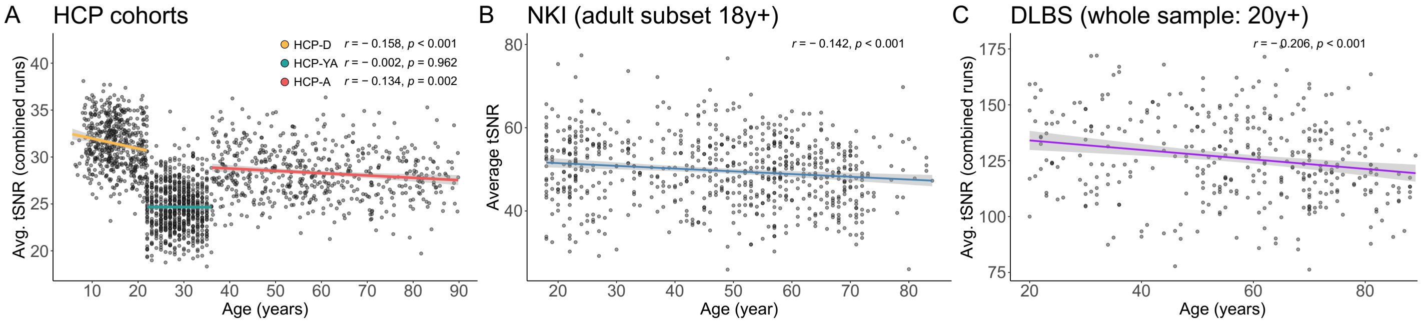

**Figure S4 – tSNR reduces with age in adulthood.**
**(A)** Age-related decreases in tSNR are observed in HCP-D (orange line), HCP-A (red line), and adults in other cohorts: **(B)** a subset of NKI samples aged 18y and older, and **(C)** the entire DLBS sample aged 20y and older.

##### *1.5 Variance and skewness of tSNR across HCP cohorts*

To evaluate whether cohort differences extended beyond shifts in average tSNR, we assessed properties of tSNR distribution across the three HCP cohorts. A median-centered Levene's test shows significant differences in variance across HCP cohorts ($F$ = 4.54, $p$ = .011), where the variance ($s^2$) of tSNR was greatest in HCP-A ($s^2$=7.57), followed by HCP-D ($s^2$=6.96) and HCP-YA ($s^2$=6.06; see **Fig. S5**). HCP-A having the largest variance was not unexpected given its wide age range (36-90y).

Using a bootstrap analysis, we compared the skewness ($g_1$) of the tSNR distribution across HCP cohorts. Skewness differed between HCP-D ($g_1$ = -0.07) and both HCP-YA ($g_1$ = 0.25, 95% CI of difference [-0.58, -0.074]) and HCP-A ($g_1$ = 0.18, 95% CI of difference [-0.50, -0.001]. The skewness of HCP-YA and HCP-A did not significantly differ from one another (95% CI of difference [-0.34, 0.19]).

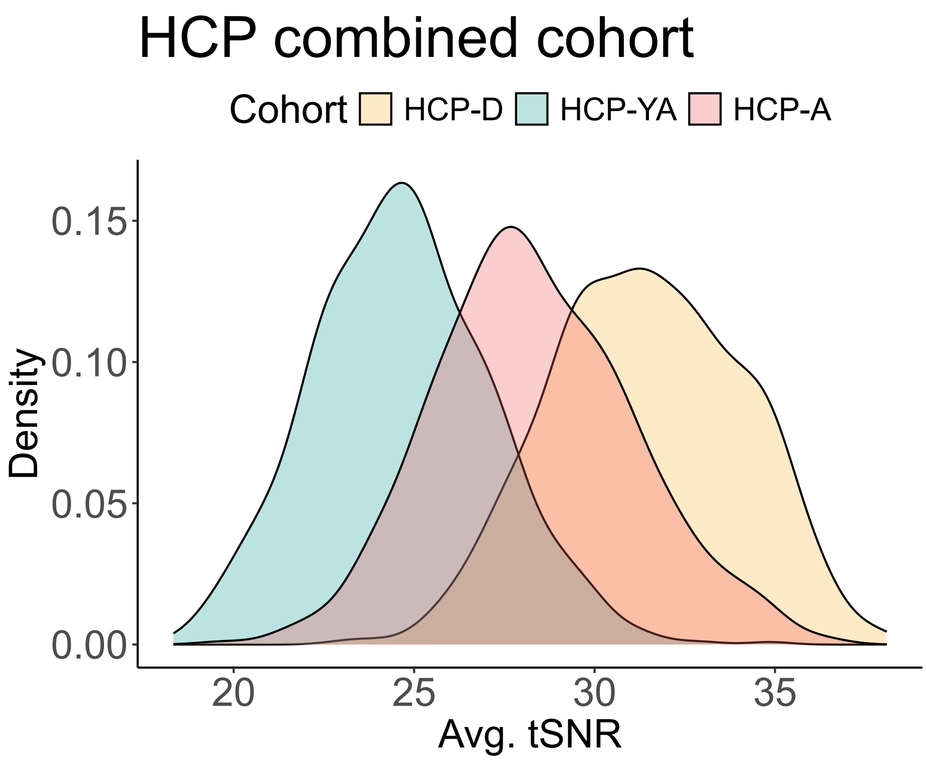

**Figure S5 – Distribution of tSNR across HCP cohorts.**
tSNR is on average higher in HCP-D, followed by HCP-A and lowest in HCP-YA. HCP-A has the largest variance, followed by HCP-D and then HCP-YA.

#### 1.6 Confidence intervals of RSFC measures

**Table S1 – Confidence intervals of RSFC measures from adjacent age intervals in HCP cohorts, independent datasets (NKI, DLBS), and harmonized HCP cohorts (CovBat and ComBat-GAM)**

| | Mean RSFC | Mean within-system RSFC | Mean between-system RSFC | System Segregation | Modularity (Q) | Mean Participation Coefficient | Mean Clustering Coefficient |
|---|---|---|---|---|---|---|---|
| **HCP cohorts before data harmonization** | | | | | | | |
| HCP-D (20-21y \| n=77) | [0.007, 0.008] | [0.186, 0.202] | [-0.012, -0.010] | [0.668, 0.686] | [0.599, 0.615] | [0.486, 0.507] | [0.232, 0.245] |
| HCP-YA (22-23y \| n=85) | [0.010, 0.013] | [0.165, 0.181] | [-0.006, -0.003] | [0.618, 0.641] | [0.555, 0.581] | [0.504, 0.532] | [0.215, 0.232] |
| HCP-YA (35-37y \| n=45) | [0.009, 0.013] | [0.143, 0.165] | [-0.005, -0.001] | [0.569, 0.610] | [0.511, 0.548] | [0.515, 0.554] | [0.196, 0.219] |
| HCP-A (36-38y \| n=29) | [0.007, 0.009] | [0.155, 0.182] | [-0.009, -0.006] | [0.625, 0.658] | [0.571, 0.594] | [0.509, 0.549] | [0.195, 0.222] |
| **Independent datasets** | | | | | | | |
| NKI (20-21y \| n= 49) | [0.003, 0.004] | [0.074, 0.090] | [-0.007, -0.005] | [0.587, 0.635] | [0.563, 0.581] | [0.423, 0.454] | [0.121, 0.135] |
| NKI (22-23y \| n= 22) | [0.002, 0.004] | [0.059, 0.086] | [-0.007, -0.004] | [0.542, 0.617] | [0.556, 0.584] | [0.436, 0.489] | [0.116, 0.139] |
| NKI (33-36y \| n= 26) | [0.001, 0.002] | [0.044, 0.063] | [-0.006, -0.003] | [0.483, 0.559] | [0.526, 0.559] | [0.441, 0.484] | [0.101, 0.119] |
| NKI (37-40y \| n = 22) | [0.001, 0.002] | [0.043, 0.065] | [-0.006, -0.003] | [0.474, 0.577] | [0.525, 0.569] | [0.417, 0.482] | [0.099, 0.119] |
| DLBS (20-21y \| n = 12) | [0.008, 0.013] | [0.204, 0.257] | [-0.015, -0.008] | [0.543, 0.605] | [0.561, 0.598] | [0.568, 0.628] | [0.306, 0.354] |
| DLBS (22-25y \| n = 13) | [0.006, 0.012] | [0.201, 0.259] | [-0.016, -0.010] | [0.553, 0.623] | [0.583, 0.622] | [0.561, 0.632] | [0.302, 0.354] |
| DLBS (33-36y \| n = 20) | [0.007, 0.011] | [0.189, 0.226] | [-0.014, -0.008] | [0.539, 0.589] | [0.564, 0.599] | [0.597, 0.642] | [0.287, 0.317] |
| DLBS (37-45y \| n = 23) | [0.006, 0.009] | [0.183, 0.227] | [-0.015, -0.009] | [0.519, 0.574] | [0.553, 0.577] | [0.612, 0.658] | [0.285, 0.315] |
| **HCP cohorts after data harmonization: CovBat** | | | | | | | |
| HCP-D (20-21y \| n=77) | [0.008, 0.009] | [0.178, 0.195] | [-0.010, -0.008] | [0.651, 0.671] | [0.591, 0.608] | [0.495, 0.517] | [0.227, 0.241] |
| HCP-YA (22-23y\| n=85) | [0.008, 0.011] | [0.177, 0.192] | [-0.009, -0.006] | [0.646, 0.666] | [0.572, 0.594] | [0.495, 0.520] | [0.218, 0.234] |
| HCP-YA (35-37y \| n=45) | [0.007, 0.011] | [0.155, 0.176] | [-0.008, -0.005] | [0.603, 0.638] | [0.530, 0.562] | [0.509, 0.544] | [0.197, 0.217] |
| HCP-A (36-38y \| n=29) | [0.008, 0.011] | [0.146, 0.175] | [-0.007, -0.004] | [0.603, 0.640] | [0.556, 0.582] | [0.514, 0.559] | [0.192, 0.220] |
| **HCP cohorts after data harmonization: ComBat-GAM** | | | | | | | |
| HCP-D (20-21y \| n=77) | [0.008, 0.010] | [0.175, 0.192] | [-0.010, -0.008] | [0.644, 0.664] | [0.579, 0.598] | [0.495, 0.519] | [0.223, 0.239] |
| HCP-YA (22-23y\| n=85) | [0.009, 0.011] | [0.179, 0.194] | [-0.009, -0.006] | [0.646, 0.667] | [0.579, 0.601] | [0.493, 0.517] | [0.224, 0.240] |
| HCP-YA (35-37y \| n=45) | [0.008, 0.011] | [0.157, 0.178] | [-0.008, -0.005] | [0.600, 0.637] | [0.538, 0.570] | [0.505, 0.539] | [0.206, 0.226] |
| HCP-A (36-38y \| n=29) | [0.008, 0.011] | [0.143, 0.172] | [-0.007, -0.004] | [0.600, 0.637] | [0.551, 0.579] | [0.518, 0.564] | [0.185, 0.215] |

The age ranges for the comparison datasets (NKI and DLBS) were extended in certain intervals to increase participant coverage. This expansion is expected to increase, rather than reduce sensitivity to age-group effects.
Color of the cell reflects statistical significance: grey = $p > .05$ *ns*, yellow = $p < .10$+, orange = $p < .05$*

#### *1.7 NKI alternative resting-state sequences' tSNR and network measures across the lifespan*

The Nathan Kline Institute Rockland Sample (NKI) dataset was used as a reference dataset. It has multiple resting-state sequences, and the one used in the main analysis was chosen due to its similarity to HCP-style data, specifically, a multiband acquisition with very short TR (645ms). The other two sequences had either a slightly longer TR of 1400ms (multiband factor of 4), or a long TR of 2500ms with no multiband.

The sequence parameters were as follows. Short-TR sequence: multiband factor=4, TR = 645ms, TE = 30ms, flip-angle = 60°, 3mm isotropic voxels, 40 slices, FOV = 200×200mm, 900 volumes (~9.7min). Mid-TR sequence: multiband factor=4, TR = 1400ms, TE = 30ms, flip-angle = 65°, 2mm isotropic voxels, 64 slices, FOV = 200×200mm, 404 volumes (~9.4min). Legacy sequence: no multiband, TR = 2500ms, TE = 30ms, flip-angle = 80°, 3mm isotropic voxels, 38 slices, FOV = 216×216mm, 120 volumes (5 min).

The lifespan age trajectories of tSNR and network measures for all three sequences were plotted in **Fig S6.** Across the three resting-state sequences, there were differences in tSNR (**Fig. S6A**) and the mean level of network measures, but the age-trajectory for all three sequences were smooth and did not show abrupt changes near the young adult age range.

Age was negatively correlated with tSNR in two out of the three sequences across the entire cohort: TR=645ms $r$ = -0.05, p=.119, TR=1400ms $r$ = -0.11, $p$ = .032, TR=2500ms $r$ = -0.11, $p$ < .001. This similar pattern persisted even when restricting to adults matching HCP-A age range (36+): TR=645ms $r$ = -0.051, p=.119, TR=1400ms $r$ = -0.11, $p$ = .032, TR=2500ms $r$ = -0.11, $p$ < .001.

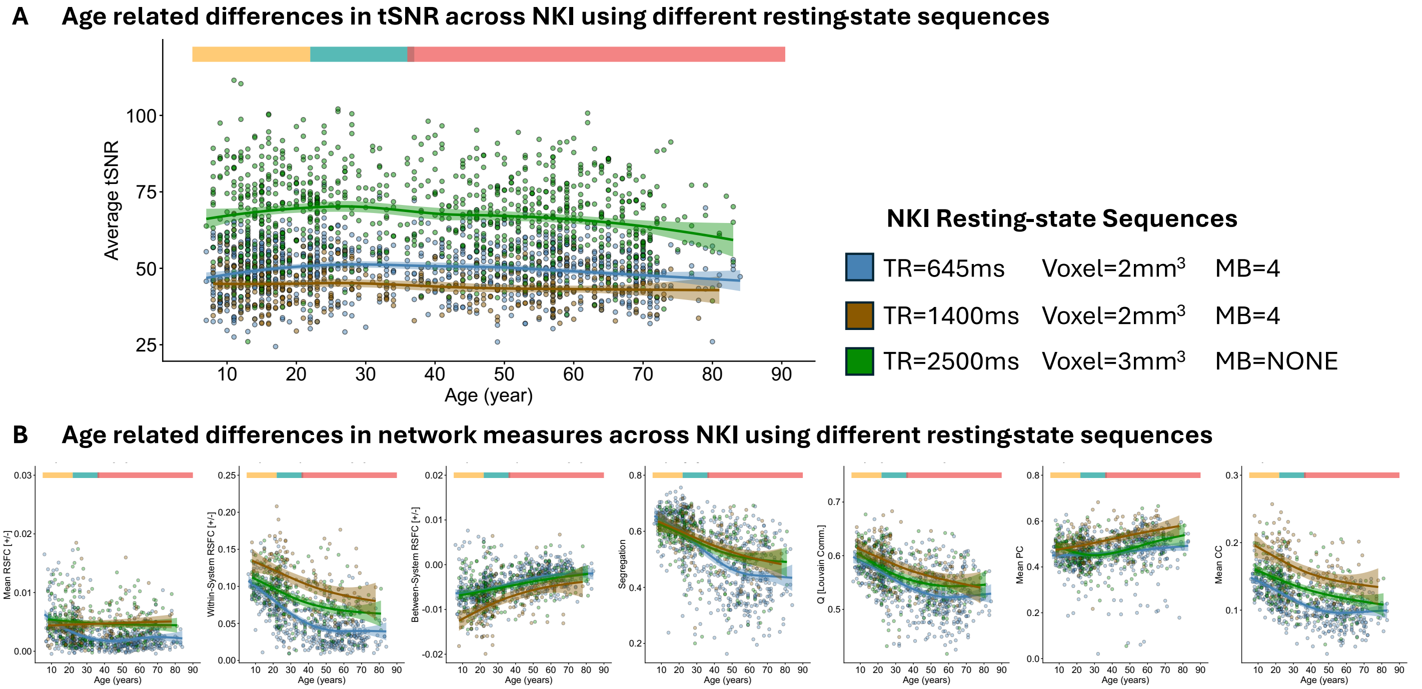

**Figure S6 – Three NKI scan protocols vary in signal characteristics, but all exhibit predictable aging trend in tSNR and network measures.**
**(A)** tSNR exhibits a smooth age trend across the NKI lifespan sample in all three scan protocols. **(B)** The seven network measures for all three resting-state protocols exhibit predictable age trend. TR = repetition time; MB = multiband.

##### *1.8 Using a more stringent data threshold (20 min) does not alter observed patterns in HCP cohorts*

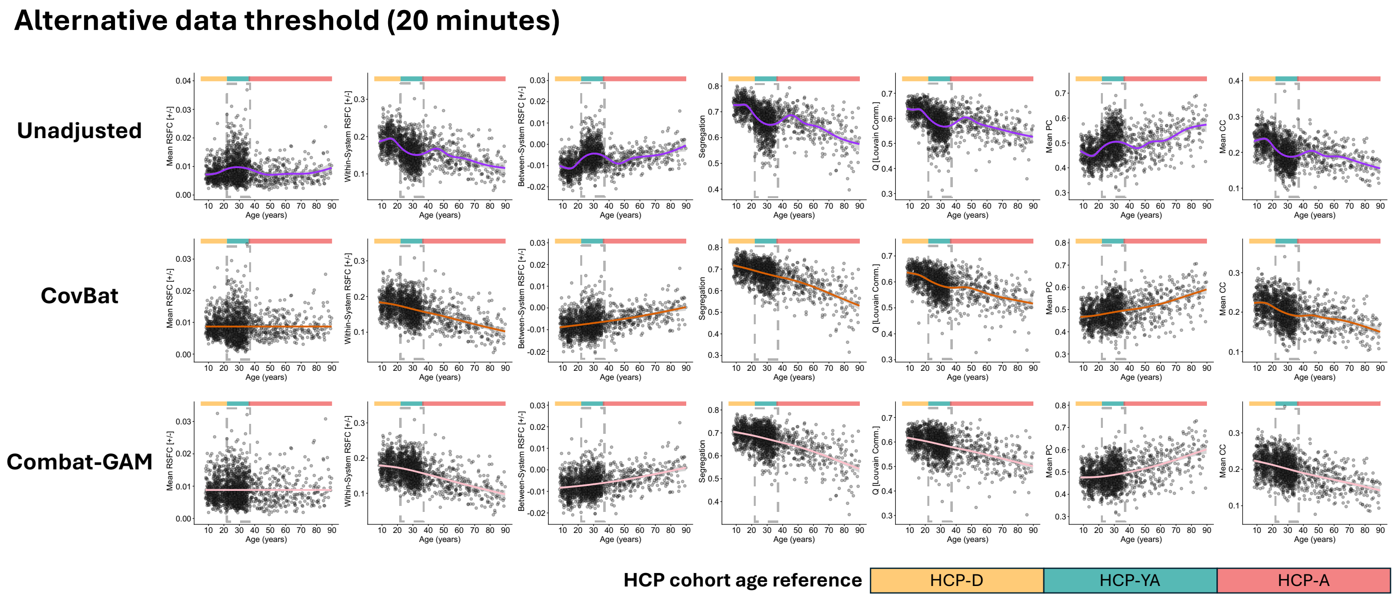

**Figure S7 – Lifespan differences in network measures across HCP cohorts using 20 minutes data threshold exhibit similar patterns to main analyses using 10 minutes data threshold.**
Generalized additive models (GAMs) were fitted to visualize age-related trends for seven measures (left to right): mean RSFC, within-system RSFC, between-system RSFC, system segregation, modularity (Q), mean participation coefficient (PC), and mean clustering coefficient (CC). Colored bars above graphs indicate the different HCP-cohorts (yellow=HCP-D, teal=HCP-YA, salmon=HCP-A) and dashed gray lines demarcate the 25-37 HCP-YA age-range for easy visualization of distinct properties of the HCP-YA data. **(A)** HCP cohorts show strong cohort-related deviations in these RSFC measures (see pink arrows). For example, the HCP-YA cohort exhibits inflated mean and between-system RSFC alongside reduced within-system RSFC and system segregation, leading to atypical age-related trajectories. These cohort-related deviations were attenuated after data harmonization using CovBat **(B)** and **(C)** ComBat-GAM.

##### *1.9 Differences in tSNR across testing sites/scanner*

HCP-YA's 3T data were all collected at Washington University (WashU) but on a different scanner than HCP-D and HCP-A data collected at WashU (custom Skyra vs Prisma). Furthermore, HCP-YA used a different acquisition protocol (resting-state fMRI TR of 720ms [LR/RL phase encoding] for HCP-YA vs. 800ms TR [AP/PA phase encoding] for HCP-D/A). The differences in scanner and acquisition were completely confounded in this case; thus, we always refer to them together (i.e., protocol/scanner model).

HCP-D/A used Siemens Prisma scanners across four different sites for each cohort (note that Harvard and MGH are two separate sites, where HCP-D participants were collected at Harvard and HCP-A participants were collected at MGH). See **Table S2** for breakdown of cohort, site, number of participants included in the present work, and their mean tSNR (SD).

Within HCP-D/A, there were site differences in tSNR. The largest site-differences observed in both HCP-D and HCP-A were between WashU and UMinn (HCP-D: $t = 7.14$, $p < .001$; HCP-A: $t = 8.83$, $p < .001$). However, tSNR in the HCP-YA cohort was lower than all the other cohort/sites (all pairwise comparison between HCP-YA and other cohort/sites were significantly different *t*s > 12.77 *p*s < .001; see **Fig. S8** for visualization of tSNR distribution across cohorts and sites).

**Table S2**
**Mean tSNR (SD) of each site within an HCP cohort**

| Cohort | Site | N | Mean tSNR (SD) |
|---|---|---|---|
| HCP-D | Harvard | 170 | 31.56 (2.47) |
| | UCLA | 107 | 31.57 (2.53) |
| | UMinn | 134 | 32.33 (2.60) |
| | WashU | 109 | 29.99 (2.49) |
| HCP-YA | WashU | 764 | 24.67 (2.46) |
| HCP-A | MGH | 112 | 27.80 (2.42) |
| | UCLA | 117 | 28.26 (2.47) |
| | UMinn | 155 | 29.84 (2.71) |
| | WashU | 157 | 27.20 (2.55) |

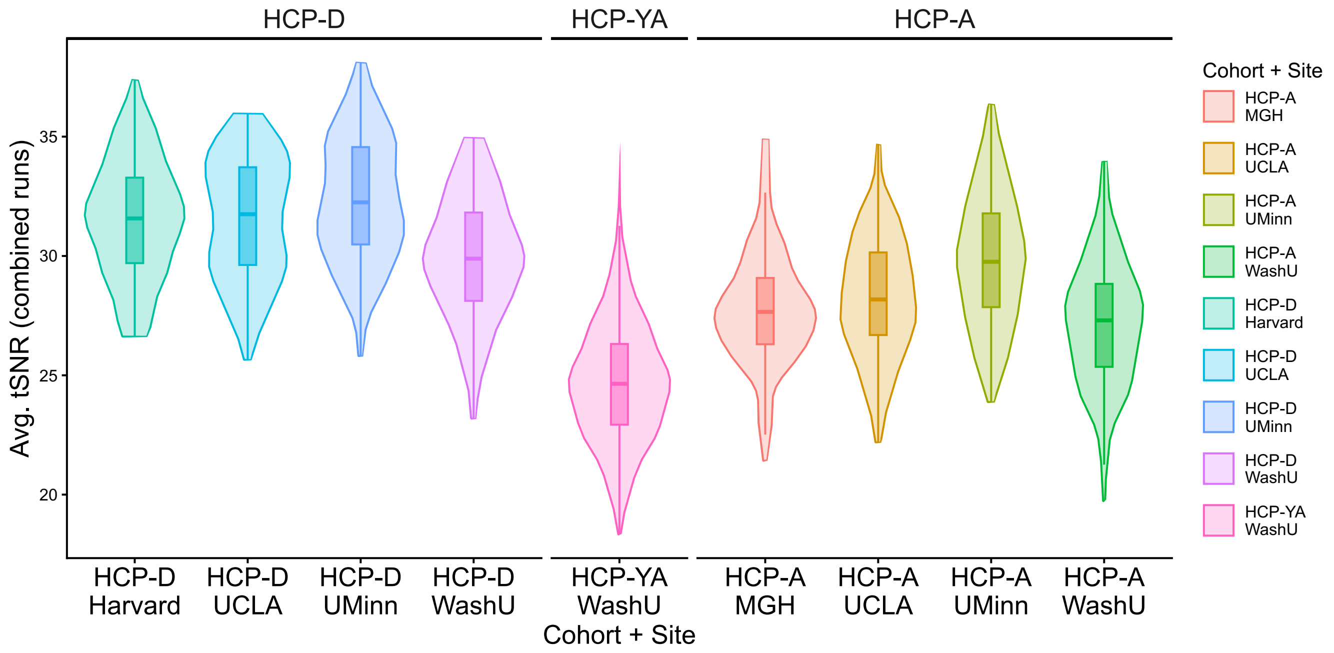

**Figure S8 – HCP signal quality as measured by tSNR across cohort and site.**
Across HCP cohorts and sites, tSNR differed within cohorts and across cohorts. However, the difference in tSNR between HCP-YA and other cohorts was greater than within-cohort site differences.

##### *1.10 Alternative harmonization*

Alternative batching strategies were used to show that data harmonization can target different effects, but most critically, in the HCP datasets, any batching strategy that isolated HCP-YA as a distinct batch yielded acceptable results.

We included four additional batching alternatives: “Cohort” separated the three datasets into their own batches (k = 3), “Site” separated the physical sites into their own batches (k = 5), “Site + Scanner Model” was similar to Site but further separated the WashU site into the data that were collected with the Prisma scanner (HCP-D & HCP-A) vs. the Skyra scanner (HCP-YA; k=6), and lastly, “Site + Cohort” separated each physical site within cohorts (k = 9). See **Table S3** for a detailed breakdown.

CovBat and ComBat-GAM were both used to target the four alternative batch effects. The network measures based on the alternative harmonization were plotted in **Fig. S9A (CovBat) and S9B (ComBat-GAM).** The only batching alternative that did not perform well was “Site” (k = 5; see **Table S4** and **Table S5**), which preserved the elevated mean RSFC and between-system RSFC in HCP-YA relative to the other two cohorts, while also retaining reduced within-system RSFC in HCP-YA. Specifically, because HCP-YA data were included in the same batch as HCP-D and HCP-A data collected at WashU, this batching strategy cannot effectively adjust for protocol and hardware differences in HCP-YA.

**Table S3 – Alternative harmonization approaches’ targeted batch effects**

| **Batching Variable** | **Protocol / Scanner Model** | **Cohort** | **Site** | **Site + Scanner Model** | **Site + Cohort** |
|---|---|---|---|---|---|
| **k (n batches)** | 2 (main) | 3 | 5 | 6 | 9 |
| **Batches** | Skyra<br>Prisma | HCP-D<br>HCP-YA<br>HCP-A | Harvard<br>MGH<br>UCLA<br>UMinn<br>WashU | Harvard-Prisma<br>MGH-Prisma<br>UCLA-Prisma<br>UMinn-Prisma<br>WashU-Prisma<br>WashU-Skyra | Harvard-HCP-D<br>MGH-HCP-A<br>UCLA-HCP-D<br>UCLA-HCP-A<br>UMinn-HCP-D<br>UMinn-HCP-A<br>WashU-HCP-D<br>WashU-HCP-YA<br>WashU-HCP-A |

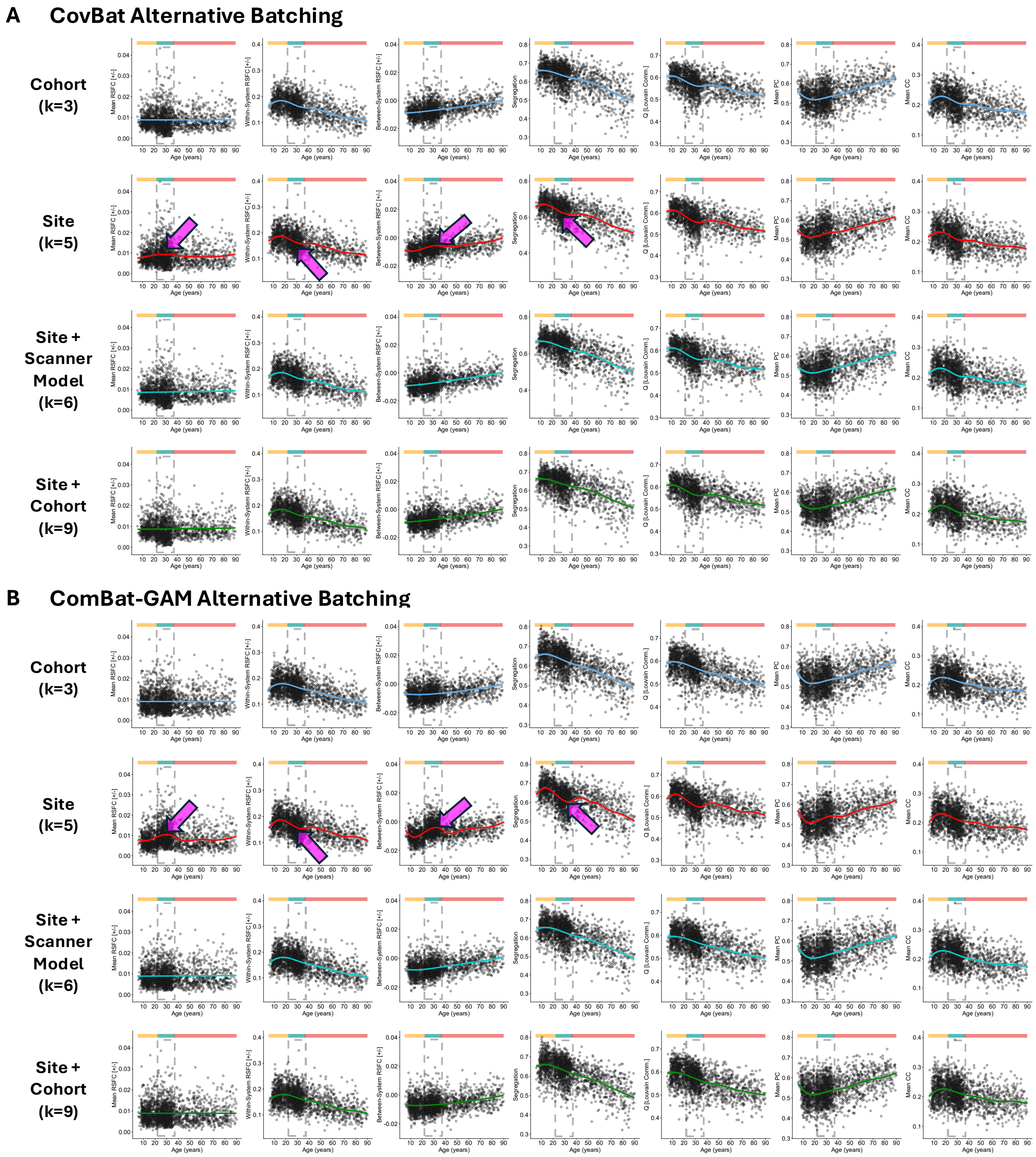

**Figure S9 – Alternative data harmonization batching using CovBat and ComBat-GAM.**
**(A)** CovBat and **(B)** ComBat-GAM were used to create harmonized network measures using four alternative batching approaches (k = number of batches). Site (k = 5) was the only batching approach that failed to remove elevated mean RSFC and between system RSFC in HCP-YA compared to the other two cohorts (see pink arrows). It also preserved the reduction in within-system RSFC among HCP-YA compared to the other two cohorts. Together, these effects resulted in a much stronger dip in system segregation when batching by Site. All other approaches that isolated HCP-YA into an individual batch yielded similar results.

**Table S4 – Comparing RSFC measures between adjacent age intervals in harmonized HCP cohorts: CovBat and ComBat-GAM with alternative batch variables**

| | | Mean RSFC | Mean within-system RSFC | Mean between-system RSFC | System Segregation | Modularity (Q) | Mean Participation Coefficient | Mean Clustering Coefficient |
|---|---|---|---|---|---|---|---|---|
| **HCP cohorts after alternate data harmonization adjusting for Cohort (k=3)** | | | | | | | | |
| **CovBat: Cohort** | HCP-D (20-21y \| n=77) vs. HCP-YA (22-23y\| n=85) | ns | ns | ns | ns | ns | ns | ns |
| | HCP-YA (35-37y \| n=45) vs. HCP-A (36-38y \| n=29) | ns | ns | ns | ns | ** | ns | ns |
| **ComBat-GAM: Cohort** | HCP-D (20-21y \| n=77) vs. HCP-YA (22-23y\| n=85) | ns | ns | ns | ns | ns | ns | ns |
| | HCP-YA (35-37y \| n=45) vs. HCP-A (36-38y \| n=29) | ns | ns | ns | ns | ns | ns | ns |
| **HCP cohorts after alternate data harmonization adjusting for Site (k=5)** | | | | | | | | |
| **CovBat: Site** | HCP-D (20-21y \| n=77) vs. HCP-YA (22-23y\| n=85) | ** | ns | ** | * | ** | ns | ns |
| | HCP-YA (35-37y \| n=45) vs. HCP-A (36-38y \| n=29) | ns | ns | ns | ns | ** | ns | ns |
| **ComBat-GAM: Site** | HCP-D (20-21y \| n=77) vs. HCP-YA (22-23y\| n=85) | *** | + | *** | *** | ** | ns | ns |
| | HCP-YA (35-37y \| n=45) vs. HCP-A (36-38y \| n=29) | ** | ns | ** | ** | *** | ns | ns |
| **HCP cohorts after alternate data harmonization adjusting for Site + Scanner Model (k=6)** | | | | | | | | |
| **CovBat: Site + Scanner Model** | HCP-D (20-21y \| n=77) vs. HCP-YA (22-23y\| n=85) | ns | ns | ns | ns | + | ns | ns |
| | HCP-YA (35-37y \| n=45) vs. HCP-A (36-38y \| n=29) | ns | ns | ns | ns | ** | ns | ns |
| **ComBat-GAM: Site + Scanner Model** | HCP-D (20-21y \| n=77) vs. HCP-YA (22-23y\| n=85) | ns | ns | ns | ns | ns | ns | ns |
| | HCP-YA (35-37y \| n=45) vs. HCP-A (36-38y \| n=29) | ns | ns | ns | ns | ns | ns | + |
| **HCP cohorts after alternate data harmonization adjusting for Site + Cohort (k=9)** | | | | | | | | |
| **CovBat: Site + Cohort** | HCP-D (20-21y \| n=77) vs. HCP-YA (22-23y\| n=85) | ns | ns | ns | ns | ns | ns | ns |
| | HCP-YA (35-37y \| n=45) vs. HCP-A (36-38y \| n=29) | ns | ns | ns | ns | ** | ns | ns |
| **ComBat-GAM: Site + Cohort** | HCP-D (20-21y \| n=77) vs. HCP-YA (22-23y\| n=85) | ns | ns | ns | ns | ns | ns | ns |
| | HCP-YA (35-37y \| n=45) vs. HCP-A (36-38y \| n=29) | ns | ns | ns | ns | ns | ns | ns |

+$p < .10$, *$p < .05$, **$p < .01$, ***$p < .001$

**Table S5 – Confidence intervals of RSFC measures from adjacent age intervals in alternatively harmonized HCP cohorts: CovBat and ComBat-GAM with alternative batch variables**

| | | Mean RSFC | Mean within-system RSFC | Mean between-system RSFC | System Segregation | Modularity (Q) | Mean Participation Coefficient | Mean Clustering Coefficient |
|---|---|---|---|---|---|---|---|---|
| **HCP cohorts after alternate data harmonization adjusting for Cohort (k=3)** | | | | | | | | |
| **CovBat: Cohort** | HCP-D (20-21y \| n=77) | [0.008, 0.010] | [0.179, 0.195] | [-0.010, -0.008] | [0.644, 0.664] | [0.578, 0.599] | [0.502, 0.524] | [0.226, 0.239] |
| | HCP-YA (22-23y\| n=85) | [0.008, 0.011] | [0.176, 0.191] | [-0.009, -0.006] | [0.645, 0.665] | [0.572, 0.594] | [0.495, 0.520] | [0.218, 0.234] |
| | HCP-YA (35-37y \| n=45) | [0.007, 0.011] | [0.155, 0.176] | [-0.008, -0.005] | [0.602, 0.637] | [0.530, 0.562] | [0.509, 0.544] | [0.197, 0.218] |
| | HCP-A (36-38y \| n=29) | [0.008, 0.010] | [0.146, 0.176] | [-0.007, -0.004] | [0.609, 0.648] | [0.566, 0.590] | [0.515, 0.558] | [0.191, 0.218] |
| **ComBat-GAM: Cohort** | HCP-D (20-21y \| n=77) | [0.008, 0.010] | [0.173, 0.190] | [-0.009, -0.007] | [0.643, 0.669] | [0.579, 0.600] | [0.499, 0.526] | [0.219, 0.235] |
| | HCP-YA (22-23y\| n=85) | [0.009, 0.011] | [0.178, 0.193] | [-0.009, -0.006] | [0.646, 0.667] | [0.579, 0.601] | [0.494, 0.518] | [0.224, 0.240] |
| | HCP-YA (35-37y \| n=45) | [0.008, 0.011] | [0.157, 0.178] | [-0.008, -0.005] | [0.600, 0.637] | [0.538, 0.570] | [0.505, 0.539] | [0.206, 0.226] |
| | HCP-A (36-38y \| n=29) | [0.008, 0.011] | [0.146, 0.174] | [-0.008, -0.005] | [0.602, 0.633] | [0.550, 0.577] | [0.513, 0.555] | [0.190, 0.220] |
| **HCP cohorts after alternate data harmonization adjusting for Site (k=5)** | | | | | | | | |
| **CovBat: Site** | HCP-D (20-21y \| n=77) | [0.007, 0.009] | [0.180, 0.196] | [-0.011, -0.009] | [0.654, 0.675] | [0.591, 0.609] | [0.494, 0.516] | [0.228, 0.241] |
| | HCP-YA (22-23y\| n=85) | [0.009, 0.012] | [0.173, 0.189] | [-0.008, -0.005] | [0.637, 0.658] | [0.567, 0.590] | [0.498, 0.524] | [0.217, 0.234] |
| | HCP-YA (35-37y \| n=45) | [0.008, 0.012] | [0.152, 0.173] | [-0.007, -0.003] | [0.593, 0.630] | [0.525, 0.558] | [0.511, 0.547] | [0.197, 0.218] |
| | HCP-A (36-38y \| n=29) | [0.007, 0.010] | [0.149, 0.178] | [-0.008, -0.005] | [0.614, 0.650] | [0.566, 0.588] | [0.513, 0.555] | [0.190, 0.218] |
| **ComBat-GAM: Site** | HCP-D (20-21y \| n=77) | [0.006, 0.008] | [0.181, 0.198] | [-0.012, -0.010] | [0.660, 0.682] | [0.592, 0.611] | [0.490, 0.515] | [0.226, 0.241] |
| | HCP-YA (22-23y\| n=85) | [0.010, 0.013] | [0.171, 0.186] | [-0.007, -0.004] | [0.627, 0.649] | [0.564, 0.587] | [0.500, 0.525] | [0.221, 0.237] |
| | HCP-YA (35-37y \| n=45) | [0.009, 0.013] | [0.149, 0.170] | [-0.006, -0.002] | [0.580, 0.618] | [0.522, 0.555] | [0.512, 0.547] | [0.203, 0.223] |
| | HCP-A (36-38y \| n=29) | [0.006, 0.009] | [0.151, 0.179] | [-0.010, -0.006] | [0.620, 0.654] | [0.569, 0.594] | [0.510, 0.552] | [0.189, 0.219] |
| **HCP cohorts after alternate data harmonization adjusting for Site + Scanner Model (k=6)** | | | | | | | | |
| **CovBat: Site + Scanner Model** | HCP-D (20-21y \| n=77) | [0.008, 0.009] | [0.179, 0.195] | [-0.010, -0.008] | [0.651, 0.671] | [0.590, 0.607] | [0.497, 0.518] | [0.226, 0.239] |
| | HCP-YA (22-23y\| n=85) | [0.008, 0.011] | [0.177, 0.192] | [-0.009, -0.007] | [0.647, 0.667] | [0.573, 0.595] | [0.495, 0.519] | [0.218, 0.234] |
| | HCP-YA (35-37y \| n=45) | [0.007, 0.011] | [0.156, 0.177] | [-0.008, -0.005] | [0.605, 0.639] | [0.531, 0.562] | [0.509, 0.543] | [0.197, 0.218] |
| | HCP-A (36-38y \| n=29) | [0.008, 0.011] | [0.147, 0.177] | [-0.007, -0.004] | [0.609, 0.646] | [0.562, 0.585] | [0.510, 0.555] | [0.191, 0.219] |
| **ComBat-GAM: Site + Scanner Model** | HCP-D (20-21y \| n=77) | [0.008, 0.010] | [0.174, 0.191] | [-0.010, -0.008] | [0.641, 0.664] | [0.578, 0.598] | [0.497, 0.522] | [0.223, 0.238] |
| | HCP-YA (22-23y\| n=85) | [0.009, 0.011] | [0.179, 0.194] | [-0.009, -0.006] | [0.647, 0.667] | [0.579, 0.601] | [0.493, 0.517] | [0.225, 0.240] |
| | HCP-YA (35-37y \| n=45) | [0.008, 0.011] | [0.157, 0.178] | [-0.008, -0.005] | [0.600, 0.637] | [0.538, 0.570] | [0.505, 0.539] | [0.206, 0.226] |
| | HCP-A (36-38y \| n=29) | [0.008, 0.011] | [0.143, 0.172] | [-0.007, -0.004] | [0.602, 0.638] | [0.553, 0.580] | [0.517, 0.561] | [0.185, 0.216] |
| **HCP cohorts after alternate data harmonization adjusting for Site + Cohort (k=9)** | | | | | | | | |
| **CovBat: Site + Cohort** | HCP-D (20-21y \| n=77) | [0.008, 0.010] | [0.180, 0.195] | [-0.010, -0.008] | [0.647, 0.666] | [0.581, 0.601] | [0.499, 0.520] | [0.225, 0.237] |
| | HCP-YA (22-23y\| n=85) | [0.008, 0.011] | [0.177, 0.192] | [-0.009, -0.007] | [0.646, 0.666] | [0.573, 0.595] | [0.495, 0.519] | [0.217, 0.234] |
| | HCP-YA (35-37y \| n=45) | [0.007, 0.011] | [0.155, 0.176] | [-0.008, -0.005] | [0.604, 0.639] | [0.532, 0.563] | [0.509, 0.544] | [0.197, 0.217] |
| | HCP-A (36-38y \| n=29) | [0.008, 0.010] | [0.146, 0.176] | [-0.008, -0.004] | [0.612, 0.650] | [0.567, 0.591] | [0.511, 0.556] | [0.188, 0.216] |
| **ComBat-GAM: Site + Cohort** | HCP-D (20-21y \| n=77) | [0.008, 0.010] | [0.173, 0.190] | [-0.009, -0.007] | [0.644, 0.670] | [0.580, 0.601] | [0.499, 0.525] | [0.219, 0.235] |
| | HCP-YA (22-23y\| n=85) | [0.009, 0.011] | [0.178, 0.193] | [-0.009, -0.006] | [0.646, 0.667] | [0.579, 0.601] | [0.494, 0.518] | [0.224, 0.239] |
| | HCP-YA (35-37y \| n=45) | [0.008, 0.011] | [0.157, 0.178] | [-0.008, -0.005] | [0.599, 0.636] | [0.538, 0.570] | [0.506, 0.539] | [0.206, 0.226] |
| | HCP-A (36-38y \| n=29) | [0.008, 0.011] | [0.145, 0.174] | [-0.008, -0.005] | [0.601, 0.633] | [0.551, 0.576] | [0.513, 0.556] | [0.189, 0.219] |

Yellow: $p < .10$; Orange: $p < .05$
